# Per- and Polyfluoroalkyl Substances Induces Salt-Sensitive Hypertension by Upregulating Epithelial Sodium Channel - The First Experimental Evidence Supporting Causality

**DOI:** 10.1101/2025.09.19.677469

**Authors:** Jing Liu, Qiongzi Qiu, Christine A. Klemens, Vineetha Garimella, Jing Huang, Nawagamuwage Lilani Dilani Perera, Carrie A. McDonough, Christina M. Post, B. Paige Lawrence, Thu H. Le, Changwei Li, Mingyu Liang, Jing O. Wu

**Affiliations:** Department of Physiology, College of Medicine – Tucson, University of Arizona, Tucson AZ; Department of Molecular Pharmacology & Physiology; Hypertension and Kidney Research Center, Morsani College of Medicine, University of South Florida, Tampa, FL; Department of Chemistry, Carnegie Mellon University, Pittsburgh, PA; Department of Environmental Medicine, University of Rochester Medical Center, Rochester, NY; Department of Medicine, University of California, Irvine, Irvine, CA; Department of Epidemiology, O’Donnell School of Public Health, University of Texas Southwest Medical Center, Dallas, TX

**Author notes:** Co-first authors (Jing Liu, Qiongzi Qiu). **Corresponding Authors**: Jing O. Wu, PhD, FAHA, Assistant Professor, Department of Physiology College of Medicine – Tucson University of Arizona Medical Research Building Rm 420, Jing Liu, PhD, Assistant Research Professor Department of Physiology College of Medicine – Tucson University of Arizona Medical Research Building 440K.

**Keywords:** Fluorochemicals, hypertension, kidney injury, salt sensitivity, epithelial sodium channel

## Abstract

Per– and polyfluoroalkyl substances (PFAS) are synthetic chemicals found in the plasma of 98% of Americans. Epidemiological studies associate PFAS exposure with hypertension and kidney dysfunction, but causality and mechanisms remain unclear. We examined the effects of a mixture of 4 PFAS commonly detected in humans, including PFOA, PFOS, PFHxS, and PFNA, on blood pressure, salt sensitivity, and renal injury in 129S6 mice. Exposure to a lower dose for 3 weeks produced plasma PFAS levels in mice resembling occupational and regional environmental exposures; while a upper dose achieved levels similar to PFAS production workers. PFAS induced dose-dependent pressor effects in male but not female mice on a 0.4% low salt diet. During 4% high salt feeding, PFAS induced greater salt-sensitive hypertension in male mice, accompanied by glomerulopathy, interstitial fibrosis, and a trend towards increased albuminuria. Pressor effects were independent of plasma norepinephrine. Single-cell RNA sequencing of kidneys revealed most transcriptional changes in proximal tubule, thick ascending limb, and collecting duct, with enrichment of pathways in cholesterol synthesis, mitochondria respiration, ATP production, and transmembrane transporter activity. PFAS markedly increased the mRNA and protein of the pore-forming α subunit of epithelial sodium channel (αENaC), with no change in βENaC and a slight reduction in γENaC protein. Elevated αENaC coincided with a 30% decrease in Nedd4-2 phosphorylation (Ser448), suggesting reduced ENaC ubiquitination and degradation. However, protein expression of α1 Na^+^-K^+^-ATPase and serum– and glucocorticoid-regulated kinase 1 (SGK1) as well as SGK1 phosphorylation (Ser78) were unaltered. Amiloride abolished salt-induced hypertension in lower-dose mice but only partially corrected hypertension in the upper dose group. Taken together, our results provide causal evidence that PFAS exposure promotes hypertension, salt sensitivity, and kidney injury via renal epithelial mechanisms, supporting and extending human epidemiologic observations.

**Translational Statement:** Our findings establish four PFAS as causal drivers of salt-sensitive hypertension and kidney injury through convergent effects on ENaC and other tubular sodium transporters. These results not only provide a mechanistic explanation for epidemiologic associations but also identify PFAS as environmental amplifiers of dietary sodium risk. Given the ubiquity of human exposure, reducing PFAS burden alongside salt reduction may represent a complementary strategy to curb the global epidemic of hypertension and kidney disease.

## INTRODUCTION

Hypertension affects 48% of US adults and is a major driver of cardiovascular and kidney diseases^1^. A key contributor to hypertension is salt sensitivity of blood pressure (SSBP), defined as >5 mmHg or 5% change of blood pressure following sodium load or depletion ^2^. SSBP affects more than 79% of African Americans and over 36% of Caucasians ^3^ and is an independent predictor of cardiovascular morbidity and mortality ^4^. Despite their high prevalence and clinical importance, the environmental factors that promote hypertension and SSBP remain poorly defined, and the molecular mechanisms are largely unknown.

Per– and polyfluoroalkyl substances (PFAS) are a large group of highly persistent synthetic chemicals detected in the blood of more than 98% of Americans ^5^. Perfluorooctanoic acid (PFOA), perfluorooctane sulfonate (PFOS), perfluorohexane sulfonate (PFHxS), and perfluorononanoic acid (PFNA) are among the most abundant PFAS in human blood and tissues ^5–7^. The exceptionally stable carbon-fluorine bonds prevent metabolism, resulting in long blood half-lives in humans (3.8 – 8.5 years) ^8,9^. Elevated blood PFAS levels are strongly associated with hypertension and kidney dysfunction in large population studies ^6,7,10,11^, but these relationships are correlative. Causality has not been established, and underlying mechanisms remain elusive.

For the general population, ingestion of PFAS-contaminated food and water is the primary exposure route^12^. PFAS bioaccumulate in the food chain ^13^ and are detectable in at least 45% US drinking water samples ^14^, leading to daily exposure to small amounts. In the National Health and Nutrition Examination Survey (NHANES) participants, median blood levels of the four PFAS above range from 1.1 to 12.8 ng/mL, reflecting exposure levels in the US general population ^6^. In contrast, some populations such as firefighters ^15^ and residents of contaminated regions^16–18^ often show blood PFAS levels 10-100 times higher than the general population. Importantly, firefighters have also higher blood pressure and increased rates of hypertension compared to civilians ^19,20^, which may be in part attributable to their chronic PFAS exposures. The most extreme exposure to date has been documented in fluorochemical production workers who have blood PFOA concentrations reaching 22,000 – 47,000 ng/mL ^21,22^, over 1000-fold above levels in the US general population, representing the upper known limit of human exposures.

Humans are exposed to a complex mixture of long-chain and short-chain PFAS, yet the combined health effects of these mixtures are poorly understood. A meta-analysis of 13 population studies with 81,096 participants found that PFOA, PFOS, PFHxS, and PFNA were independently associated with hypertension, while other types of PFAS (ΣPFAS, perfluorodecanoic acid, perfluoroundecanoic acid) had no statistical significance ^23^. To model real world exposure, we developed an oral drinking-water protocol containing a quaternary mixture of the human hypertension-relevant PFOA, PFOS, PFHxS, and PFNA in a 1: 1: 0.2: 0.14 mass ratio, matching their relative abundance in human blood ^24^. Mice were exposed to either an upper dose PFAS mixture or a lower dose PFAS mixture. The upper dose corresponds to the highest occupational exposures in fluorochemical production workers ^21,22,24^, while the lower dose (1:100 dilution of the upper dose) represents chronic low level occupational or regional environmental exposures.

Because urinary excretion is a major route of PFAS elimination, especially for shorter chain perfluorocarboxylates^25,26^, the kidney is a key target for their toxic effects. The primary objective of this study was to directly test whether oral PFAS exposure promotes hypertension, SSBP, and kidney injury in mice, and to elucidate renal tubular mechanisms that may drive these effects. Using an integrated approach combining analytical chemistry, cardiovascular monitoring, single-cell RNA sequencing (scRNAseq), molecular biology, and *in vivo* pharmacology, we identified a mechanistic link between PFAS exposure, renal tubular sodium reabsorption, and elevated blood pressure. This study provides experimental evidence that supports a causal relationship between PFAS exposure and hypertension. These findings not only advance our mechanistic understanding but also have direct implications for PFAS regulation, environmental policy, and the prevention of exposure-related cardiovascular risk.

## METHODS

### Animals and treatments

All protocols were approved by the Animal Care and Use Committees at the University of Rochester and the University of Arizona. Mouse husbandry and care in the study followed the National Institutes of Health (NIH) guidelines. The 129S6 mice were purchased from Taconic Biosciences (129S6/SvEvTac). Mice were maintained on a 0.4% low salt diet (Dyet #113755 AIN-76A purified rodent diet) upon arrival until they were provided with the 4% high salt diet (Dyet #113756 AIN-76A purified rodent diet). Each cage was supplied with environmental enrichments, and the animal facility was maintained on standard 12-hour light-dark cycles. Tap water supplied by the animal housing facilities was provided to mice until they received special water containing the lower and upper concentrations of the PFAS mixture. Mice were randomized into control, lower PFAS dose, or upper PFAS dose groups using a random number sequence generated with the RANDBETWEEN function in Excel. Each mouse was assigned a random number, sorted in ascending order, and then allocated sequentially to the three groups.

The upper dose, consisting of 0.009 mg/mL PFOA + 0.009 mg/mL PFOS + 0.0018 mg/mL PFHxS, and 0.0013 mg/mL PFNA in drinking water ^24^, which delivered an estimated daily dose of 1.88 mg/kg PFOA, 1.88 mg/kg PFOS, 0.376 mg/kg PFHxS, and 0.26 mg/kg PFNA (Table 1A). The lower dose group received water containing 1:100 dilution of the upper dose, which is estimated to deliver 18.8 μg/kg/day PFOA, 18.8 μg/kg/day PFOS, 3.76 μg/kg/day PFHxS, and 2.6 μg/kg/day PFNA. These estimated daily doses were based on our prior observations that a 20-gram mouse typically consumed 4 mL water/day on a chow diet or a low salt diet ^27^. Because water consumption doubled on the 4% high salt diet ^27^, the concentrations of PFAS added to the water were reduced by 50% in the high salt phase to maintain the same exposure levels. Some mice received amiloride at 20 mg/kg/day in drinking water after salt-induced hypertension was established. Blood pressure and heart rate were chronically monitored using radiotelemetry as previously described ^27–29^.

**Table 1.** Oral PFAS Exposure and the Resultant Plasma Concentrations in Mice. **A**) Mice were exposed to upper and lower doses of a quaternary PFAS mixture containing PFOA, PFOS, PFHxS, and PFNA, the top 4 PFAS species detected in human blood as described in a recent publication by Post CM *et al* ^24^. The upper dose mimics occupational exposures such as fluorochemical workers; whereas the lower dose (1:100 dilution of the upper dose) mimics chronic exposures in firefighters and residents near chemical plants. Dose calculation was based on daily water consumption of 4 mL for a 20-gram mouse. **B**) Male controls received tap water provided by the institutional animal housing facility, while male experimental mice consumed drinking water containing the lower or upper PFAS doses for 21 days. Plasma PFAS concentrations in male mice in the present study were compared against circulating levels of female mice in the Post CM *et al* study since both studies included the upper dose following identical exposure protocols and PFAS levels were measured by the same researcher with liquid chromatography with quadrupole-time-of-flight mass spectrometry. Detection limit (DL) was 0.054 ng/mL for PFOS (linear + branched) and 1.59 ng/mL for PFNA. Data were expressed as mean ± SEM. * p<0.05 vs control; # p<0.05 upper dose vs lower dose; † p<0.05 male upper dose vs female upper dose. Because standard deviations were significantly different among groups, data were analyzed with Brown-Forsythe ANOVA and Kruskal-Wallis test. Male: control (n=9), lower dose (n=11), upper dose (n=10); Female: upper dose (n=3).

|  | Mass Ratio | Upper Dose<br>(mg/kg/day) | Dilution<br>Factor | Lower Dose<br>(µg/kg/day) |
| --- | --- | --- | --- | --- |
| **PFOA** | 1 | 1.88 | 1:100 Dilution<br>→ | 18.8 |
| **PFOS** | 1 | 1.88 |  | 18.8 |
| **PFHxS** | 0.2 | 0.376 |  | 3.76 |
| **PFNA** | 0.14 | 0.26 |  | 2.6 |

**Table 1 Oral PFAS Exposure and the Resultant Plasma Concentrations in Mice. A)**
|  | Plasma Concentrations (ng/mL) After 3-week Exposure |  |  |  |
| --- | --- | --- | --- | --- |
|  | Male mice in the present study |  |  | Female mice in the<br>Post *et al* study <sup>24</sup> |
|  | Control | Lower Dose | Upper Dose | Upper Dose |
| **PFOA** | 5.5 ± 3.7 | 1466 ± 357 * | 56475 ± 5566 * # † | 22313 ± 5229 |
| **PFOS** | <DL | 664 ± 143 | 36163 ± 4606 * # | 31886 ± 2488 |
| **PFHxS** | 1.4 ± 0.9 | 708 ± 144 * | 24772 ± 1684 * # † | 1083 ± 116 |
| **PFNA** | <DL | 115 ± 29 | 6326 ± 1112 * # | 7149 ± 1079 |

### Real-time quantitative PCR and Western blot

Male 129S6 mice at 8 weeks of age without radiotelemetry implantation received plain tap water or water containing the lower or upper PFAS doses on low salt for 3 weeks then on high salt for another week. After sacrifice, kidney cortex and medulla were separated under dissecting microscope, snap-frozen in liquid nitrogen, and stored in –80 freezer. Real-time quantitative PCR and western blot were performed following published protocols shown in Supplemental Methods. A full list of antibodies used is shown in Supplemental Table 1.

### Additional methods

including metabolic cage studies, urinary albumin quantification of PFAS by liquid chromatography with quadrupole-time-of-flight mass spectrometry (LC-QTOF-MS), Masson’s trichrome staining, and single-cell RNA Sequencing (scRNAseq) are described in detail in Supplemental Material.

### Statistics

All results were expressed as mean ± SEM. GraphPad Prism (version 10.4) was used for statistical analyses. Time course data of blood pressure and heart rate were analyzed by two-way ANOVA with repeated measurements. Group means were analyzed with one-way ANOVA. Tukey’s multiple comparison tests were performed for pairwise comparisons. When standard deviations were significantly different among groups (Brown-Forsythe test and Bartlett’s test), the data were analyzed with the Brown-Forsythe ANOVA or Welch ANOVA test. For data not exhibiting Gaussian distribution, Kruskal-Willis test was performed. A p-value <0.05 was considered statistically significant.

## RESULTS

### PFAS dosing produces human-relevant exposures

At 8 weeks of age, male 129S6 mice received plain tap water or drinking water containing the lower or upper dose of a quaternary PFAS mixture (Table 1A) for three weeks. Plasma PFAS concentrations, measured by LC-QTOF-MS as described ^24,30^, confirmed human-relevant exposures. Control mice displayed trace PFOA at 5.5 ± 3.7 ng/mL and PFHxS at 1.4 ± 0.9 ng/mL in the plasma (Table 1B), comparable to background levels in the NHANES participants (PFOA median 3.3 ng/mL with an interquartile range of 2.1 – 5.1 ng/mL; PFHxS median 1.7 ng/mL with an interquartile range of 0.9 – 2.9 ng/mL)^6^. PFOS and PFNA were below detection limits in control mice. The lower PFAS dose increased plasma concentrations to 1,466 ± 357 ng/mL (PFOA), 664 ± 143 ng/mL (PFOS), 708 ± 144 ng/mL (PFHxS), and 115 ± 29 ng/mL (PFNA), representing the blood levels observed in occupationally exposed firefighters (PFOS: 92 – 343 ng/mL, PFHxS: 49 – 326 ng/mL) ^15^ and residents near chemical plants (PFOA: 71 ± 151 ng/mL)^17^.

In a prior study, female mice receiving the upper PFAS dose in drinking water for three weeks exhibited circulating concentrations ranging from 1,000 to 30,000 ng/mL ^24^. To determine if this dose achieves similar effects in both sexes, we treated male 129S6 mice to the same upper PFAS dose. Blood samples in both studies were collected through similar procedures and PFAS quantification was performed by the same researcher. The levels of PFOS and PFNA were comparable between sexes (Table 1B). Male mice in the present study exhibited markedly greater levels of PFOA and PFHxS compared to that of female mice in the Post *et al* study ^24^. Importantly, both male and female mice exposed to the upper PFAS dose displayed high PFOA levels in the circulation (male 56,475 ± 5,566 ng/mL; female 22,313 ± 5,529 ng/mL). These are comparable to that of some fluorochemical production workers (up to 22,000 – 47,000 ng/mL) ^21,22^, representing the upper known limit of human exposures.

### PFAS raises blood pressure in males but not female mice on low salt

To determine the pressor effects of chronic PFAS exposures, 8-week-old male and female 129S6 mice were implanted with radiotelemeters for continuous cardiovascular monitoring (Figure 1A). After a 12-day recovery, blood pressure was recorded for 3 days at baseline. Mice were then randomly divided into three groups, which received either plain tap water (Ctrl), the lower, or upper PFAS dose for 3 weeks (day 0 – day 21). All mice consumed a 0.4% low salt diet. There was no change in systolic blood pressure (SBP) in control mice, but PFAS exposure induced time-dependent increases of SBP in male but not female mice (Figure 1B, 1E). To illustrate pressor effects of PFAS, SBP averages in week 3 were plotted against baseline values. Both PFAS doses significantly raised SBP in male but not female mice (Figure 1C and 1F). PFAS-induced change of SBP were dose-dependent in male but not female mice (ΔSBP in mmHg: Figure 1D, Male Ctrl 1.0±2.7, Lower 4.9±2.5, Upper 6.2±2.1; Figure 1G Female Ctrl 2.1±2.1, Lower 2.6±4.8, Upper 0.2±1.8).

**Figure 1.**
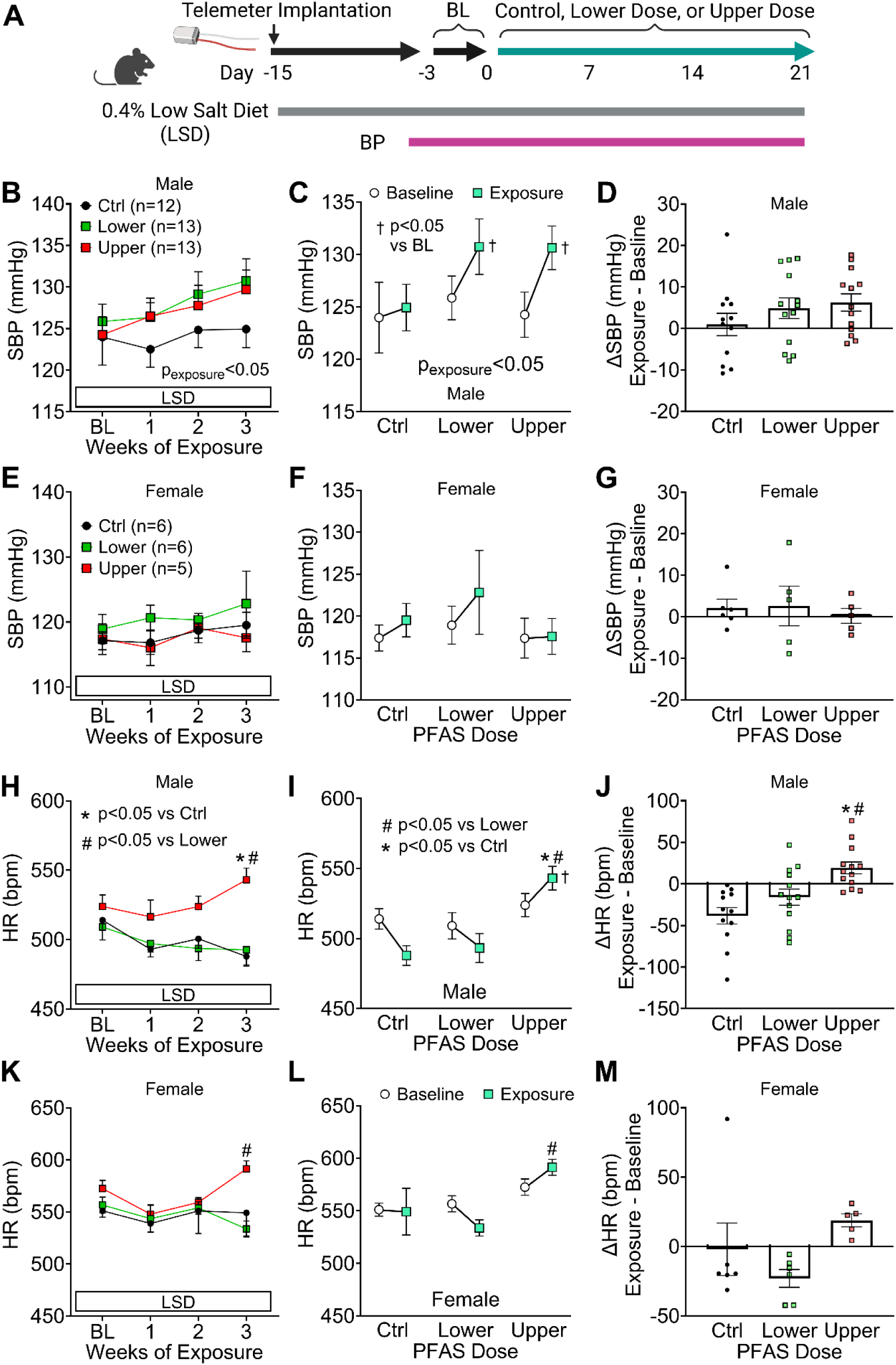
Sex-Specific Pressor Effects of PFAS on a Low Salt Diet. **A**) Study protocol. Eight-week-old male and female mice were implanted with radiotelemetry and rested for 12 days. After a 3-day baseline (BL), control mice received tap water provided by the animal facility (Ctrl), while experimental mice received water containing PFAS lower dose or upper dose for 3 weeks (day 0 – day 21). All mice consumed a 0.4% low salt diet (LSD). Systolic blood pressure (SBP) and heart rate (HR) are shown as mean ± SEM. **B-G**) Weekly SBP (B, E), before-after comparisons (C, F), and changes of SBP over 3 weeks (D, G). **H-M**) Weekly HR (H, K), before-after comparisons (I, L), and changes of HR over 3 weeks (J, M). Time course data were analyzed by 2-way ANOVA with repeated measurements, while ΔSBP and ΔHR were analyzed with 1-way ANOVA. For male mice SBP (**B**), p_exposure_<0.05, p_dose_>0.05, p_interaction_>0.05; and ΔSBP (**D**), p=0.30. For female mice SBP (**E**), p_exposure_>0.05, p_dose_>0.05, p_interaction_>0.05; and ΔSBP (**G**), p=0.85. For male mice HR (**H**), p_exposure_<0.05, p_dose_<0.01, p_interaction_<0.01; and ΔHR (**J**) p<0.01. For female mice HR (**K**), p_exposure_=0.10, p_dose_=0.23, p_interaction_<0.05; and ΔHR (**M**), p=0.10. * p<0.05 Upper vs Ctrl; # p<0.05 Upper vs Lower; †, P<0.05 Exposure vs Baseline, Tukey’s multiple comparison tests. Male n = 12, 13, 13; Female n = 6, 6, 5.

Compared to male mice that received control water or water containing the lower PFAS dose, male mice receiving the upper PFAS dose had significantly higher heart rate over the 3-week exposure (Figure 1H, 1I). The change in heart rate (ΔHR) was significantly different among three groups (Figure 1J, Ctrl –38 ± 9.9 beats per minute; lower dose –16 ± 9.8 bpm; upper dose 19 ± 7.2 bpm, 1-way ANOVA p<0.05). Compared to males, female mice exhibited similar but less pronounced changes in heart rate (Figure 1K-L) and ΔHR was not statistically different among 3 groups (Figure 1M, Ctrl –2 ± 19 bpm; lower dose – 23 ± 6.4 bpm; upper dose 19 ± 4.7 bpm, p=0.10). In both sexes, blood pressure and heart rate exhibited normal day-night rhythm, and PFAS-induced increases occurred in both the light and dark phases (Supplemental Figure 1).

### PFAS augments SSBP in a sex-dependent manner

To determine the effect of PFAS exposure on SSBP, mice were switched to a 4% high salt diet (HSD) on day 21 – day 28 while they continued to drink control or PFAS water (Figure 2A). In response to the 1-week high salt challenge, there was mild but significant BP elevation in control male 129S6 mice (Figure 2B, 2D), consistent with previous report of salt sensitivity in this strain ^31,32^. PFAS-exposed male mice exhibited greater responses to salt loading compared to male controls (Figure 2D, SBP in mmHg, Ctrl 133±2, Lower Dose 139±4, Upper Dose 140±2). In comparison, salt did not raise BP in female mice receiving control water or the lower PFAS dose, and only mildly increased SBP in the upper PFAS dose group (Figure 2C, 2E, Ctrl 121±2, Lower Dose 123±4, Upper Dose 125±3). The changes of SBP between the low salt phase and high salt phase are shown in Figure 2F. PFAS dose-dependently augmented salt-induced BP elevation in both sexes (p_exposure_<0.05) with greater SSBP in male mice than females (p_sex_<0.01). In male mice, salt loading further increased heart rate in the control and upper PFAS dose groups; but there was no effect in female mice (Figure 2G-2J). The salt-induced changes in heart rate were not different between sexes or among the dose groups (Figure 2K). During the high salt challenge, the circadian rhythm of blood pressure and heart rate was also reserved (Supplemental Figure 2).

**Figure 2.**
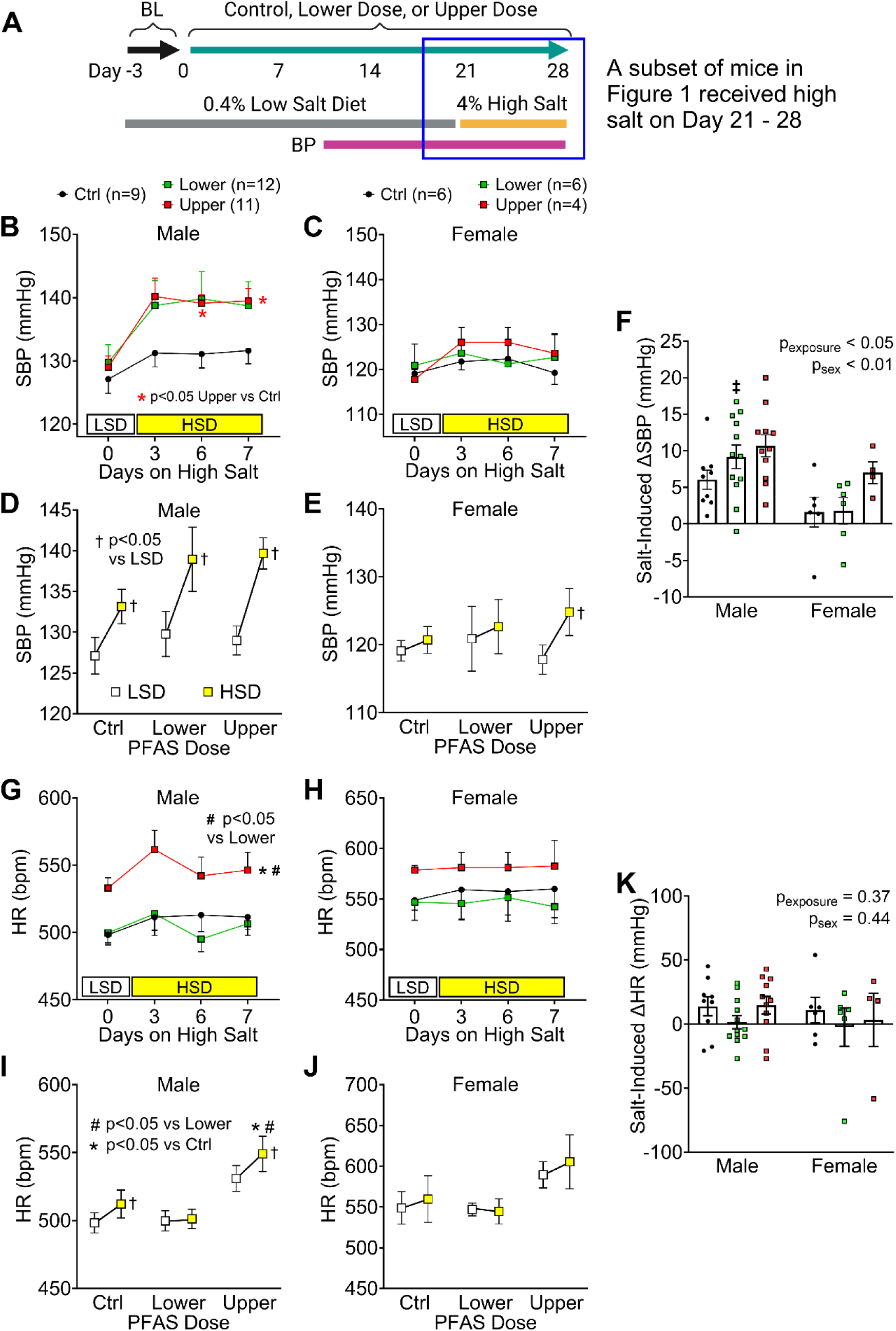
PFAS Augments Salt-Sensitive Hypertension in a Sex-Specific Manner. **A**) Protocol to assess the salt sensitivity of blood pressure. After 3-week low salt diet (LSD), a subset of mice in Figure 1 were switched to a 4% high salt diet (HSD) on day 21 – 28 (*i.e.* HSD day 1 – day 7). **B-C**) SBP responses at the end of LSD and on day 3, 6, and 7 of the 1-week HSD in male mice (the time frame in the blue box). Male: p_exposure_=0.19, p_diet_<0.01, p_interaction_=0.06; Female: p_exposure_>0.05, p_diet_<0.05, p_interaction_>0.05, 2-way ANOVA repeated measures. * p<0.05 Upper Dose vs Ctrl. **D-E**) Average SBP at the end of the low salt phase and during the 1-week high salt challenge in male mice. † p<0.05 HSD vs LSD. 2-way ANOVA repeated measures. **F**) Salt-induced SBP change in male mice. ‡ p<0.05, Male vs Female, 2-way ANOVA Tukey’s multiple comparison tests. **G-H**) Heart rate (HR) responses at day 3, 6, 7 of the 1-week HSD in female mice. Male: p_exposure_<0.01, p_diet_>0.05, p_interaction_>0.05; Female: p_exposure_>0.05, p_diet_>0.05, p_interaction_>0.05. Tukey’s multiple comparison tests, * p<0.05 Upper Dose vs Ctrl, # p<0.05 Upper Dose vs Lower Dose. **I-J**) Average SBP at the end of the low salt phase and during the 1-week high salt challenge in female mice. **K**) Salt-induced SBP change in female mice. Male, n = 9, 12, 11; Female, n = 6, 6, 4.

At the end of 3-week low salt and 1-week high salt, male but not female in the upper PFAS dose group compared to controls exhibited decreased body weight and significantly increased heart weight/body weight ratio, consistent with the hypertension phenotype (Supplemental Table 2). Consistent with prior reports on PFAS-induced spleen atrophy ^33,34^, there were sex-independent decreases in spleen weight/body weight ratio in the upper dose group. Liver to body weight ratio and kidney to body weight ratio also increased in both sexes, partly driven by the decreased body weight in the upper dose group. In comparison, there was no change in body weight or organ weight in the lower dose group.

### Pressor effects of PFAS not associated with changes in circulating norepinephrine

To determine if the effects of PFAS on blood pressure and heart rate were associated with increased sympathetic nerve activity, we measured norepinephrine levels in the plasma at the end of the 1-week high salt diet. There was no change in circulating norepinephrine levels (Supplemental Figure 3), arguing against a role of sympathoexcitation in PFAS-induced hypertension and tachycardia.

### PFAS exposure induces greater kidney damage in male mice

Glomerulosclerosis, interstitial fibrosis, and albuminuria are hallmarks of hypertensive kidney damage ^35–37^. To determine the deleterious effects of PFAS exposure in kidney pathology, we performed Masson’s trichrome staining in the kidney after the 1-week high salt diet. Male mice treated with lower and upper PFAS doses exhibited mesangial hypercellularity (Figure 3A, yellow arrows). Moreover, PFAS dose-dependently induced peri-glomerular and peri-tubular fibrosis when both sexes were analyzed together (two-way ANOVA p_exposure_<0.05, Figure 3B). In post-hoc analysis, the upper PFAS dose significantly increased renal fibrosis in PFAS-exposed male mice, which was associated with a trend toward increased urinary albumin (Figure 3C). In comparison, there was no significant kidney remodeling or albuminuria in PFAS-exposed female mice. Because female mice were largely resistant to the deleterious effects of PFAS, subsequent studies focused on male mice.

**Figure 3.**
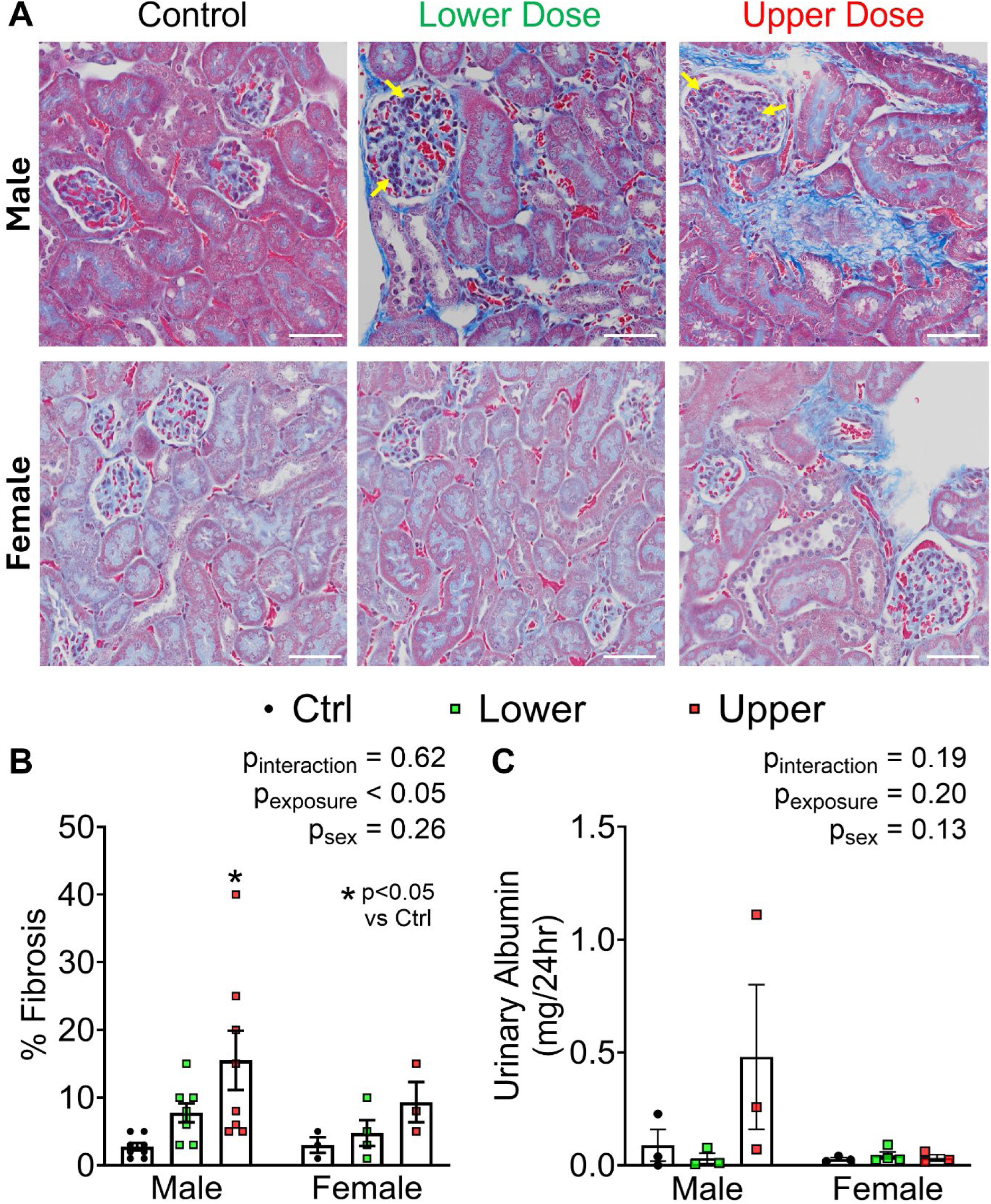
PFAS Exposure Induces Greater Kidney Damage in Male Mice. **A**) Kidney fibrosis assessed by Masson’s trichrome staining performed after 4-week exposure in a subset of mice in Figure 2. Scale bar = 60 µm. **B**) Renal fibrosis was quantified by a blinded technician experienced with renal pathology. **C**) Urinary albumin excretion was measured in 24-hour urine collected in a subset of mice from Figure 2 at the end of 1-week high salt. Data were shown as mean ± SEM and analyzed by 2-way ANOVA. Male n=8; female n=3-4, * p<0.05 Upper Dose vs Control in Tukey’s multiple comparison tests.

### Mechanistic insights from single-cell transcriptomics

To explore molecular pathways underlying PFAS-induced hypertension and SSBP at single-cell resolution, we performed single-cell RNA sequencing (scRNAseq) on whole kidneys from a separate cohort of male mice consuming control or PFAS-containing water while on a 0.4% low-salt diet for three weeks (Figure 4A). Using annotation markers in Figure 4B, we identified cell populations from the glomeruli, renal tubules, vasculature, and immune system as shown in the UMAP plots (Figure 4C). There was no change in the abundance of renal parenchymal cells, and the apparently greater composition of renal epithelial cell populations was primarily caused by reduced renal immune cell abundance in response to PFAS, especially the upper dose (Figure 4D). The decreased renal immune cell composition was consistent with decreased spleen weight in the upper dose group, reflecting PFAS-induced spleen atrophy and hypocellularity in lymphoid organs ^33,34^. Differentially expressed gene (DEG) analysis of renal tubular cells revealed that PFAS exposure induced the greatest transcriptional changes in proximal tubule (PT), thick ascending limb 2 (TAL-2), and collecting duct (CD) cells (Figure 4E). Of note, a small PT cell population was identified on the UMAP plot as a distinct cluster, with substantially more DEGs than PT S1-S3 segment cells. Compared with controls, the lower PFAS dose altered the expression of 2,707 genes in PT, 1,329 in TAL-2, and 771 in CD cells; the upper PFAS dose altered 2,041, 1,742, and 1,473 genes in these cell types, respectively (Figure 4E). Overall, the upper dose resulted in greater number of DEGs across cell types compared to the lower dose (p<0.05, one-tailed paired Wilcoxon test).

**Figure 4.**
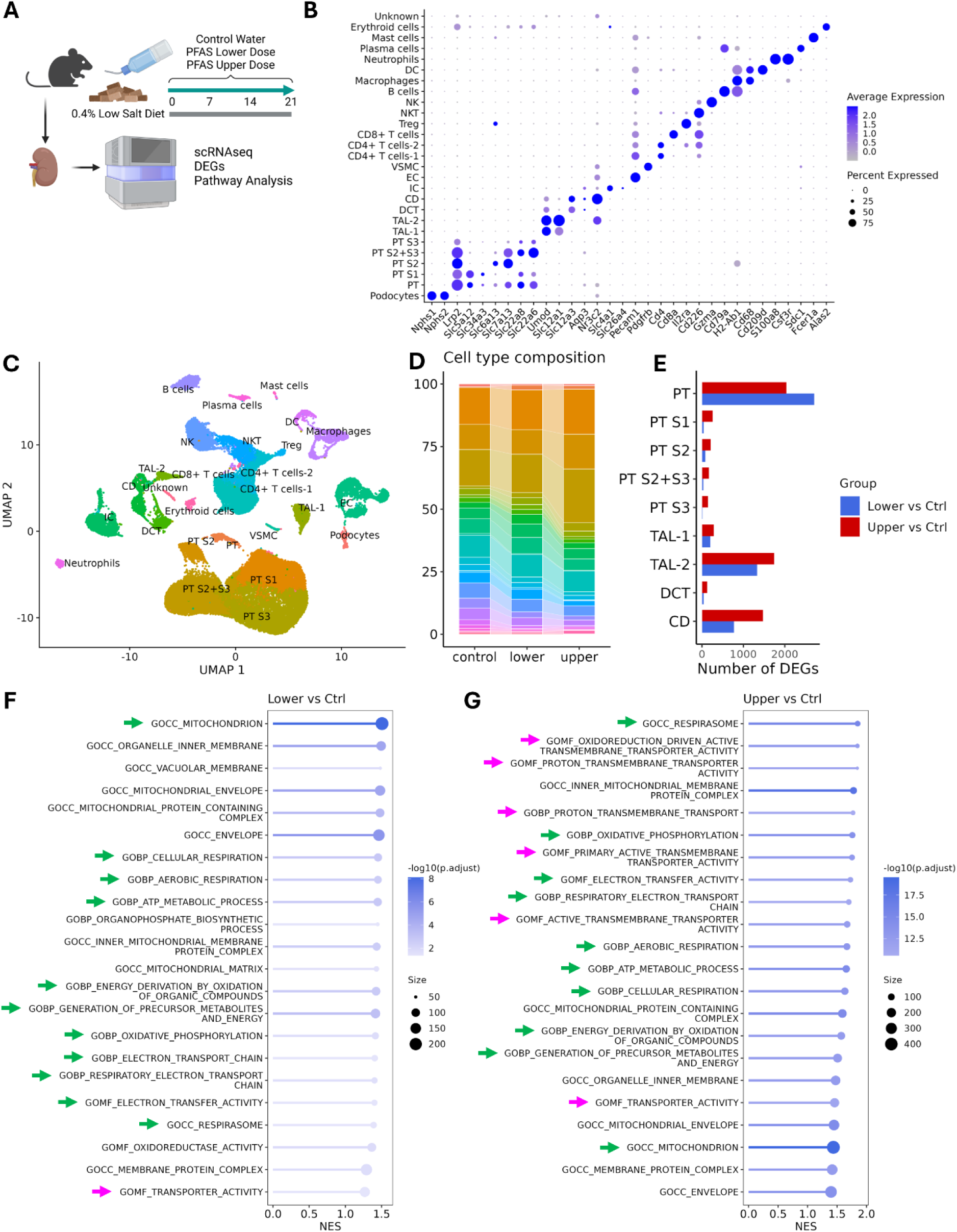
Single-cell RNA Sequencing Identifies Altered Pathways Involved in Cellular Metabolism and Transmembrane Transporter Activity. **A**) Schematic overview of scRNAseq data generation from kidneys of male 129S6 mice after treatment of control or PFAS-containing water on a 0.4% low salt diet for 3 weeks. N=3 mice per group. **B)** Dot plot showing cell type-specific expression of curated marker genes. **C)** UMAP visualization of all cells, colored and labeled by annotated cell types. **D)** Stacked bar plot showing the relative proportion of each cell type across treatment groups. **E)** Bar plot showing the number of differentially expressed genes (DEGs) identified in each cell type between lower dose vs. control and upper dose vs. control comparisons. Overall, the upper dose provoked more DEGs than the lower dose, p<0.05, one-tailed paired Wilcoxon test. **F-G)** Pathway enrichment analysis of DEGs from the collecting duct between the lower dose (F) and upper dose (G) groups compared to controls. Green arrows highlight pathways upregulated in fatty acid metabolism, mitochondria electron transfer chain, oxidative phosphorylation, and energy production. Magenta arrows highlight pathways involved in active transmembrane transporter activities.

Pathway enrichment analysis showed that the upper PFAS dose upregulated pathways in lipid metabolism, mitochondrial electron transport chain, oxidative phosphorylation, and ATP synthesis in the proximal tubule (Supplemental Figure 4A-4D), thick ascending limb (Supplemental Figure 5), and collecting duct cells (Figure 4F-4G, green arrows). The upper PFAS dose significantly increased the expression of genes involved in cholesterol influx and synthesis in the proximal tubule, including low density lipoprotein receptor (*Ldlr*), sterol regulatory element-binding protein-2 (*Srebf2*), 3-hydroxy-3-methylglutaryl-CoA (HMG-CoA) synthase 2 (*Hmgcs2*), and HMG-CoA reductase (*Hmgcr*)^38,39^ (Supplemental Figure 4G-4J). The lower PFAS dose also enriched pathways involved in mitochondria respiration and ATP synthesis only in proximal tubule S2 cells, not in S1 or S3 segments (Supplemental Figure 4E). Interestingly, the small PT cell population distinct from S1-S3 segment cells (Figure 4C) exhibited downregulated mitochondrial metabolic pathways in response to the lower PFAS dose, possibly representing injured cells (Supplemental Figure 4F, blue arrows). Collectively, these data suggest that there may be increased energy demand and expenditure as well as cholesterol synthesis in the renal epithelium during PFAS exposure.

### Expression of renal sodium transporters in response to PFAS

There were also enriched pathways in transmembrane transport activity in the proximal tubule, thick ascending limb and collecting duct cells (Supplemental Figure 4A-4E, Supplemental Figure 5, Figure 4F-G, magenta arrows), suggesting that PFAS may enhance renal tubular sodium transport. To assess the expression of key sodium transporters, we analyzed their transcription in single-cell populations and measured their mRNA levels in kidneys of additional mice by quantitative real-time PCR (rtPCR). Sodium-hydrogen exchanger 3 (NHE3, *Slc9a3*), the primary sodium transporter in the proximal tubule, was upregulated by the lower PFAS dose in scRNAseq, but this change was not detected in rtPCR (Figure 5A). Instead, rtPCR revealed a modest reduction in NHE3 mRNA with the upper PFAS dose. Other sodium transport proteins including sodium glucose co-transporter 2 (SGLT2, *Slc5a2*) and sodium phosphate co-transporter 2 (Npt2a, *Slc34a1*) also contribute to sodium reabsorption in the proximal tubule. In scRNAseq analysis, SGLT2 was upregulated in PT-S1 cells by the upper PFAS dose, while Npt2a was upregulated in PT, PT-S1, and PT-S2 cells by both PFAS doses (Supplemental Figure 4K-4L).

**Figure 5.**
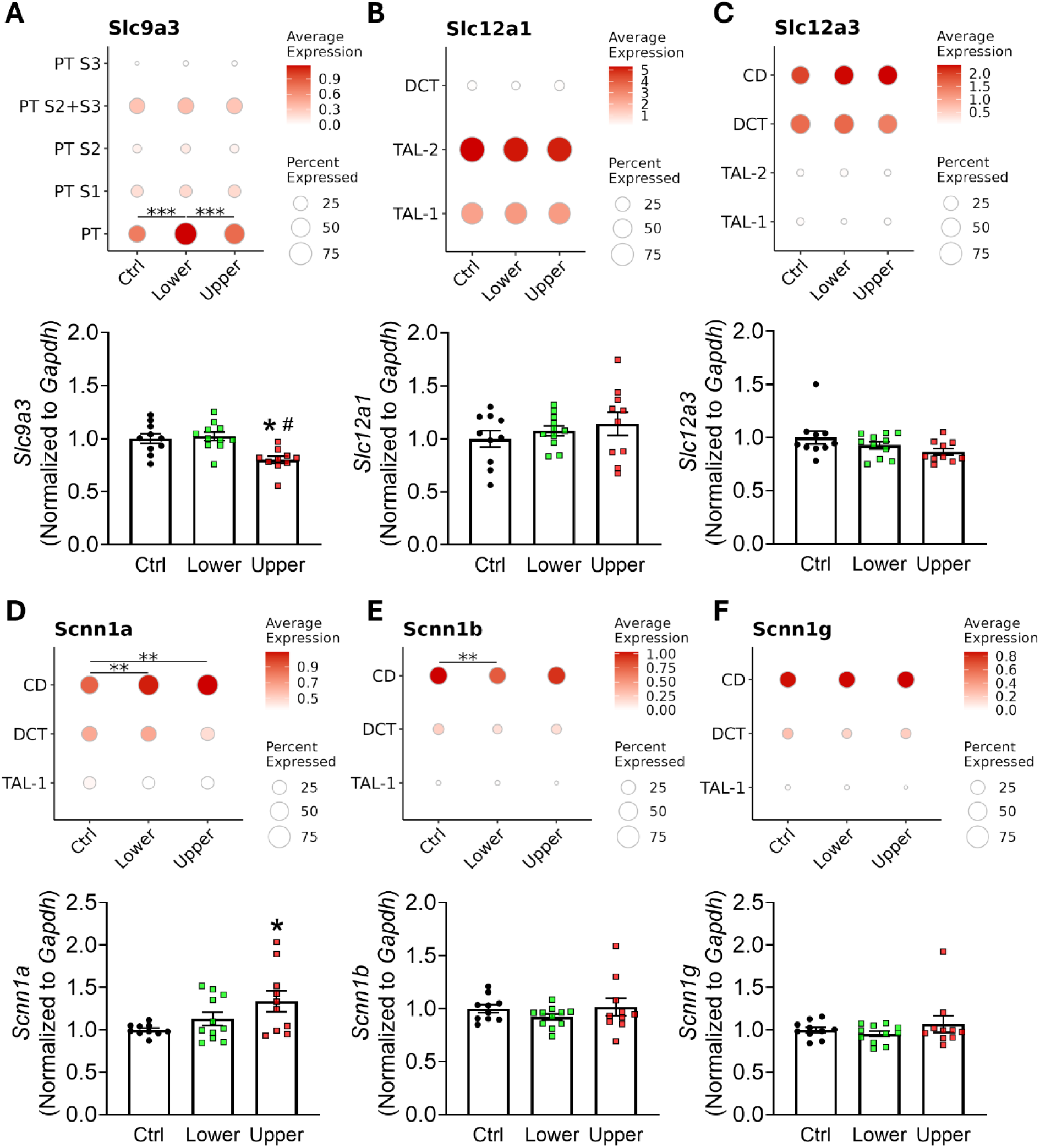
PFAS-induced Transcriptional Changes in Renal Tubular Sodium Transporters. **A-E**) Dot plots illustrate cell type-specific expression of *Slc9a3*, *Slc12a1*, *Slc12a3*, *Scnn1a*, *Scnn1b*, and *Scnn1g* in renal epithelial cells identified in the scRNAseq analysis. The size of the dots symbolizes the percentage of cells expressing genes of interest, while the depth of red color represents the levels of gene expression. For intra-cell-type comparisons across experimental groups, DEGs were computed using the Wilcoxon rank-sum test using the FindMarkers function in Seurat. Adjusted p-values were indicated for between-group comparisons on dot plots, * p<0.05, ** p<0.01, *** p<0.001. **Bar graphs** demonstrate the mRNA expression of renal sodium transporters determined by quantitative real-time PCR in the kidney cortex (*Slc9a3*, *Slc12a3*, *Scnn1a, Scnn1b & Scnn1g*) and medulla (*Slc12a1*) of additional male mice fed 3-week low salt diet + 1-week high salt diet. Data were expressed as mean ± SEM and were analyzed with 1-way ANOVA with Tukey’s multiple comparisons. For real-time PCR data, * p<0.05 upper dose vs control; # p<0.05 upper dose vs lower dose.

Sodium-potassium-2 chloride co-transporter (NKCC2, *Slc12a1*) and sodium-chloride co-transporter (NCC, *Slc12a3*) expression was unaltered in either analysis (Figure 5B-C). Consistent with this, western blotting showed no change in total NCC protein. However, there was a trend towards increased NCC phosphorylation at the Thr53 residue (pNCC, 1-way ANOVA p=0.17) and a 50% increase in pNCC/NCC ratio in the lower PFAS dose group (Supplemental Figure 6).

In collecting duct cells, epithelial sodium channel α (αENaC, *Scnn1a*), the pore-forming subunit, was significantly upregulated by both lower and upper PFAS doses in scRNAseq, consistent with a ∼40% increase in *Scnn1a* mRNA in response to the upper PFAS dose in rt-PCR (Figure 5D). PFAS-induced upregulation of αENaC was confirmed at the protein level in western blot (Figure 6A-6B). In contrast, although there was a slight decrease of βENaC (*Scnn1b*) in the lower PFAS dose group in scRNAseq and a mild reduction of γENaC (*Scnn1g*) protein in the upper PFAS dose group, there were no consistent changes in these two subunits across scRNAseq, rt-PCR, and western blot (Figure 5E-5F, and Figure 6C-6D).

**Figure 6.**
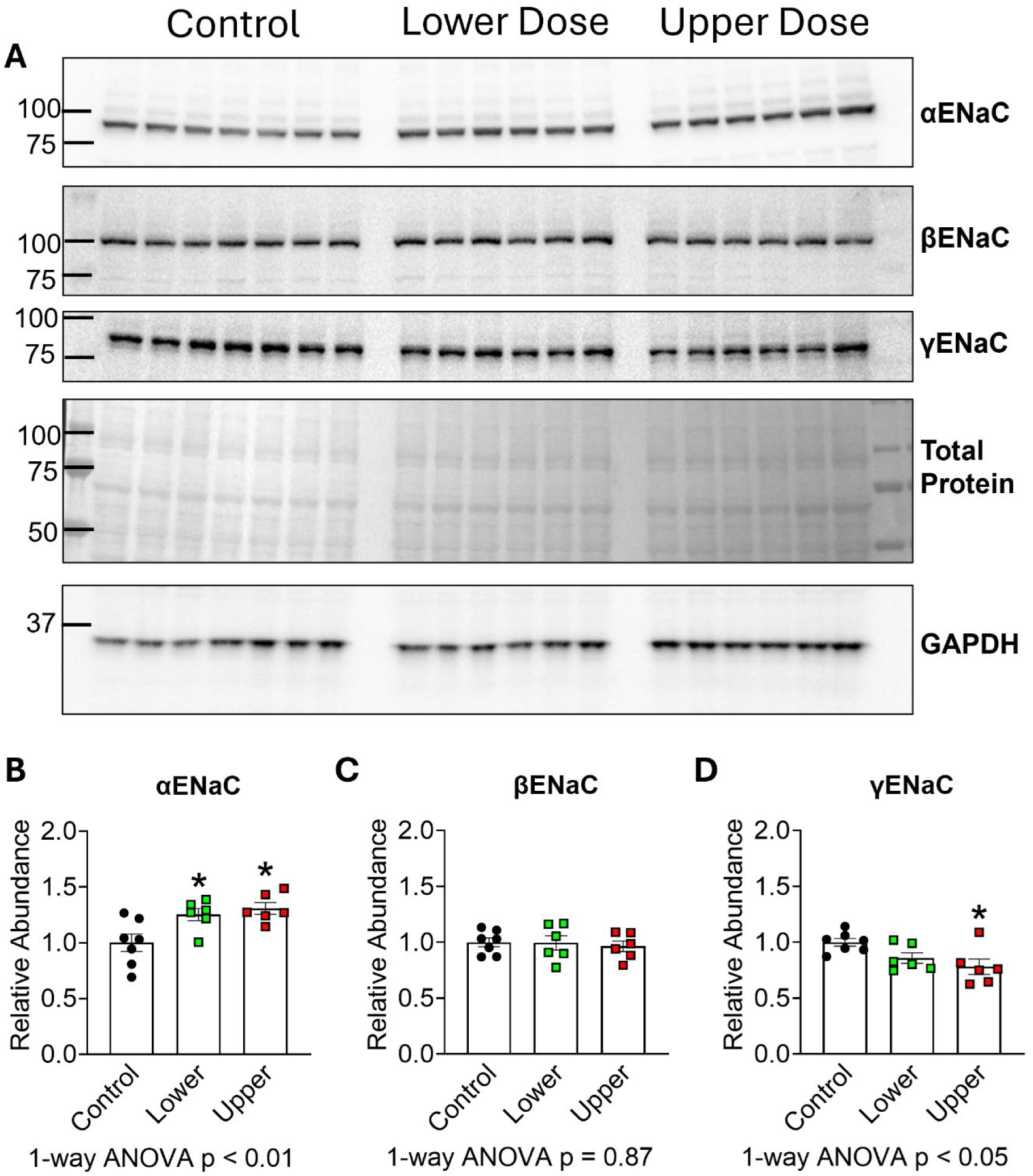
ENaC Expression in Control and PFAS-exposed Mice Fed High Salt. Control and PFAS-exposed male 129S6 mice received low salt diet for three weeks followed by 4% high salt diet for a week. Kidney cortex homogenates were used for western blot analysis of the α-, β-, and γ-ENaC subunits. Only full length α– and γ-ENaC were measured as cleaved forms could not be detected in all groups. The decrease in full length γ-ENaC may be due to increased cleavage. Each lane/dot is an individual mouse; N = 7 control, 6 lower dose, and 6 upper dose. Data are mean ± SEM. One-way ANOVA with Dunnett’s post hoc analysis. * p < 0.05 vs control.

To identify potential drivers of ENaC upregulation, we examined Nedd4-2, a ubiquitin ligase that regulates ENaC endocytosis and degradation ^40–42^. There was 12% increase of Nedd4-2 protein but a 30% decrease in Nedd4-2 phosphorylation at the S488 residue in response to the upper PFAS dose, resulting in markedly decreased pNedd4-2/Nedd4-2 ratio (Figure 7A-D). There was no change in total or pNedd4-2 in response to the lower PFAS dose. Neither total or phosphorylated (S78) serum/glucocorticoid-regulated kinase 1 (SGK1), which modulates ENaC trafficking and activity via phosphorylation of Nedd4-2 and direct interaction with ENaC ^40,41^, were unchanged; nor was there alternation in α1 sodium-potassium ATPase (α1-NKA), which transports sodium and potassium across the basolateral membrane of kidney tubules creating driving force for Na^+^ entry via ENaC (Figure 7A, 7E-G).

**Figure 7.**
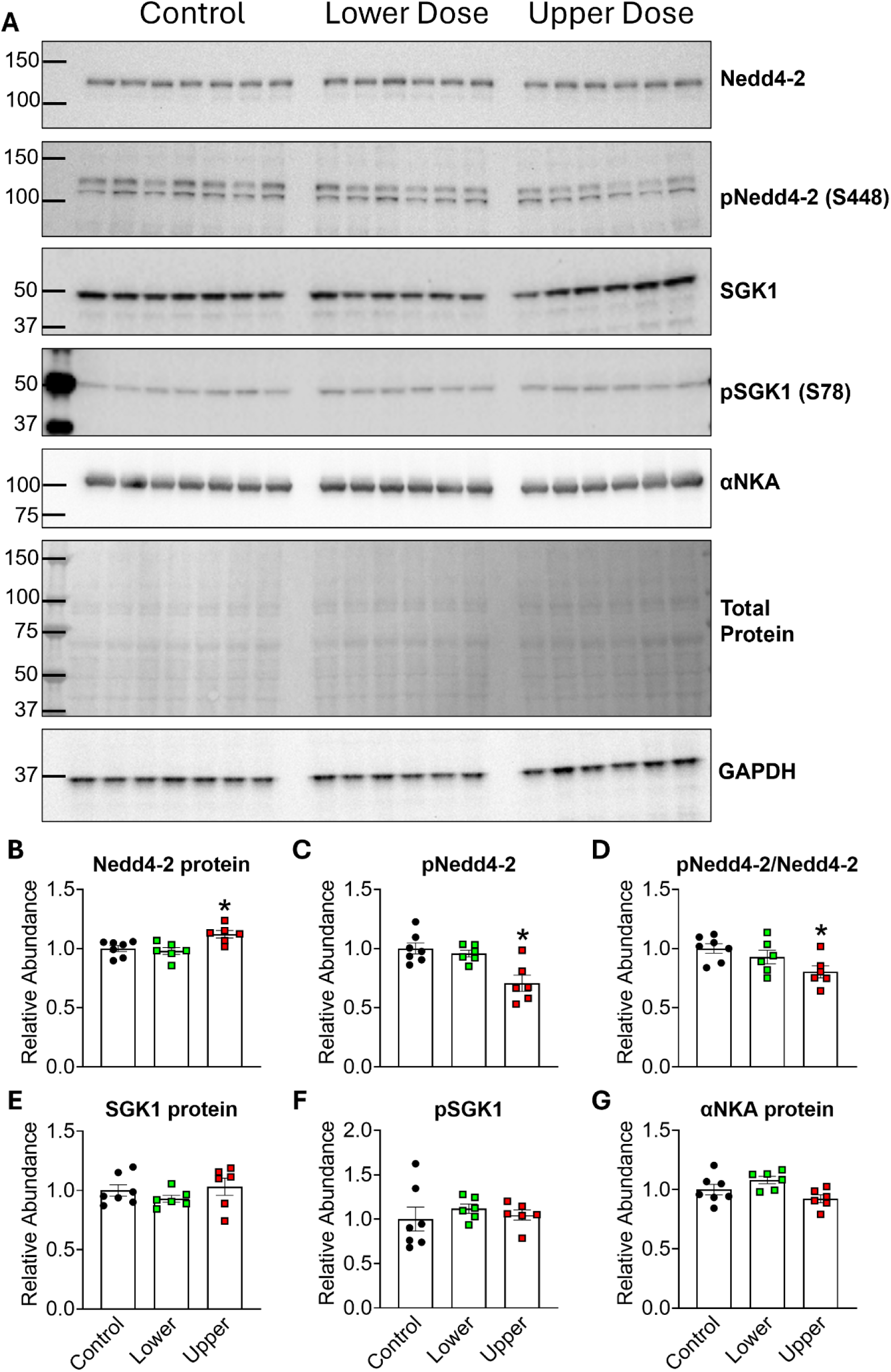
Effects of PFAS Exposure on ENaC-associated Proteins. Control and PFAS-exposed male 129S6 mice received the 0.4% low salt diet for 3 weeks followed by the 4% high salt diet for a week and kidney cortex homogenates were used for western blot. **A)** Western blot analysis of p/Nedd4-2, p/SGK1, and α1-Na^+^/K^+^ATPase (αNKA). Each lane/dot is an individual mouse; N = 7 vehicle, 6 lower dose, and 6 upper dose. **B-G**) Densitometry data of Nedd4-2, phosphorylated Nedd4-2 (Ser448), pNedd4-2/total Nedd4-2 ratio, SGK1, phosphorylated SGK1 (Ser78), and α1-NKA. Data are mean ± SEM. One-way ANOVA p<0.05 for panels B, C, and D. * p < 0.05 vs control in Dunnett’s post hoc analysis.

To assess the functional importance of ENaC in PFAS-induced salt sensitive hypertension, a subset of mice from Figure 2 was maintained on the high salt diet for an additional two weeks (day 21-35). After salt-induced hypertension was established in the first week of salt loading (day 21-28), mice received amiloride in drinking water (20 mg/kg/day, day 28-35; Figure 8A). Average SBP values before and after amiloride treatment were compiled into 24-hour plots to illustrate the depressor effects (Figure 8B-D). In the first week of high salt feeding, SBP increased by 5.6 ± 1.1 mmHg in controls, 10.8 ± 2.0 mmHg in the PFAS lower dose group, and 10.6 ± 2.7 mmHg in the PFAS upper dose group (Figure 8E). Amiloride reduced SBP by 7.3 ± 1.4 mmHg in controls and by 12.2 ± 1.9 mmHg in the PFAS lower dose group, completely abolishing the salt-induced increase. In contrast, amiloride lowered SBP by 5.4 ± 1.2 mmHg in the PFAS upper dose group, only partially reversing the salt-induced elevation. These findings indicate that ENaC mediates the exaggerated salt sensitive hypertension induced by the lower PFAS dose, whereas the pressor effects of the upper dose likely involve additional mechanisms.

**Figure 8.**
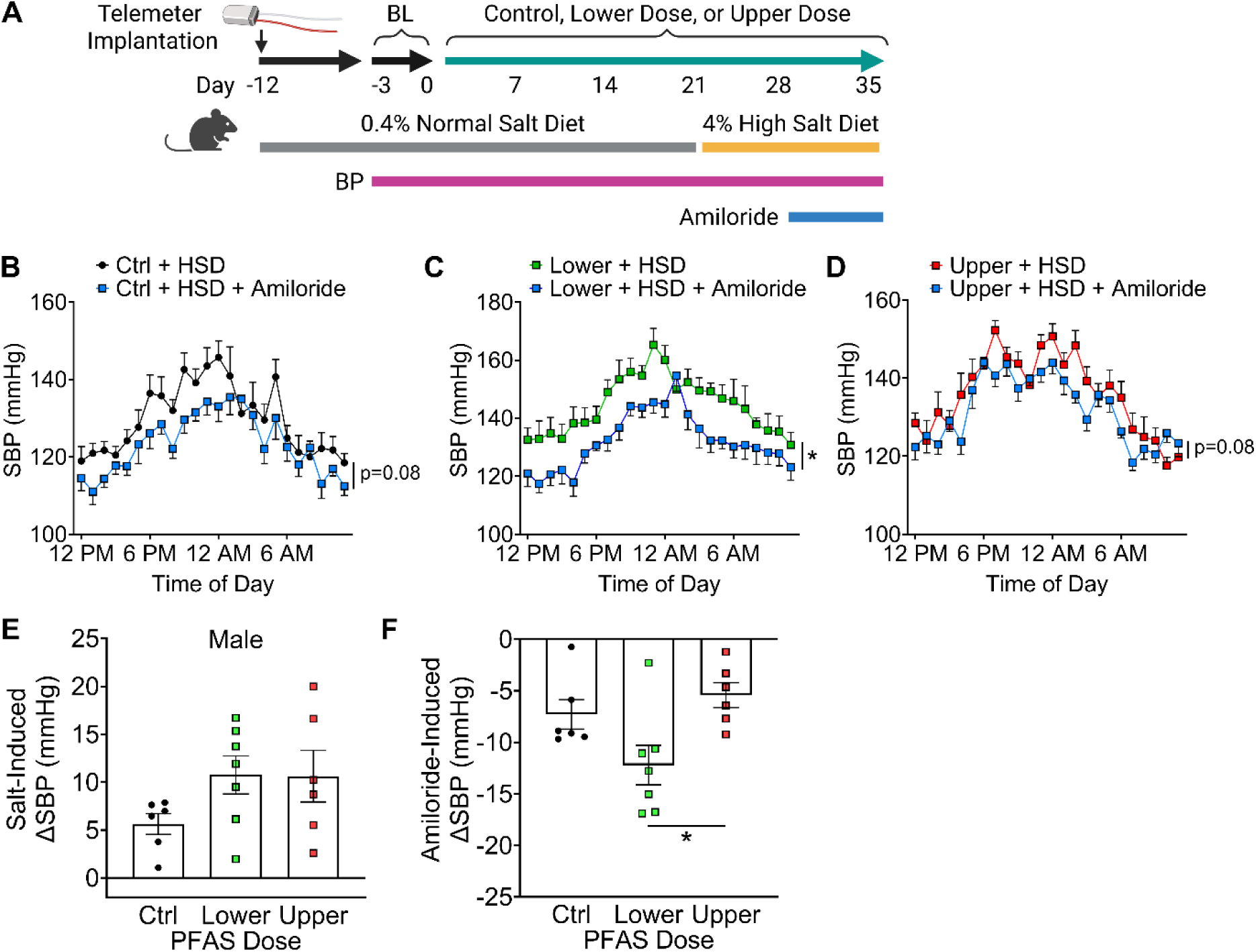
Functional Evidence Supporting a Role of ENaC in PFAS-induced Salt-Sensitive Hypertension. **A**) A subset of male mice in Figure 2 consumed the high salt diet for two weeks (day 21-35) and received amiloride at 20 mg/kg/day in drinking water during the second week of high salt feeding (day 28-35). **B-D**) The hourly blood pressure after the first week of high salt was plotted against the blood pressure after amiloride treatment. For control, lower dose and upper dose groups, n = 6, 7, 6. Data were shown as mean ± SEM and analyzed with two-way ANOVA repeated measurements. * p<0.05, effect of amiloride. **E**) Salt induced SBP elevation in the first week of high salt. **F**) Amiloride-induced reduction of blood pressure. * p<0.05, 1-way ANOVA.

## DISCUSSION

PFAS are persistent environmental contaminants linked epidemiologically to cardiometabolic, renal, hepatic, and neurocognitive conditions ^43^. While population studies consistently associate PFAS exposure with hypertension and kidney dysfunction, causal evidence has been lacking ^6,7,10,11^. Here, we provide direct experimental evidence that chronic PFAS exposure induces hypertension and kidney injury in mice, supporting a causal relationship and advancing a new paradigm in PFAS toxicology.

The public health implications are substantial. PFAS are detected in the blood of 98% of the US population ^5^, who has an average daily sodium intake (3.3 g) far exceeding the federal recommendation of 2.3 g/day ^44^. Our findings that PFAS exposure elevates blood pressure during low-salt intake and increases SSBP suggest that PFAS may contribute to the high prevalence of hypertension, particularly in the context of excessive dietary sodium intake. While sodium reduction interventions can lower blood pressure in clinical studies, population-scale salt restriction efforts have been largely ineffective due to widespread processed and restaurant foods, industrial resistance, consumer preferences, and lack of awareness ^45^. Thus, reducing PFAS exposure may represent a more tractable intervention. Recent US environmental protection agency drinking water standards are a critical first step ^46^, but further action is needed to limit dietary PFAS exposure in the general population, reduce occupational exposure in high-risk groups, and remediate environment contamination.

The PFAS doses tested in the present study approximate blood levels observed in chronically exposed groups, including fluorochemical production workers ^21,22^, firefighters ^15^ and residents exposed to contaminated water supplies ^17^, populations numbering in the millions in the US alone and much more worldwide. Both doses elevated blood pressure and enhanced SSBP, but the absence of a significant difference between the lower and the upper dose suggests a plateau effect. This raises the possibility that even exposures below these levels, including background exposures in the general population, may be sufficient to elevate blood pressure. Supporting this, NHANES analyses show that blood PFOA concentrations >1.8 ng/mL are associated with 85% higher odds ratio of hypertension per 10-fold increase, and PFNA >0.53 ng/mL with a 64% increase ^6^. Because human exposure spans a continuum from background to occupational levels, additional studies are warranted to determine the pressor effects of chronic low-level exposure.

Sex differences are another important dimension. Men consistently exhibit higher blood PFAS concentrations than women, both in the US general population ^5^ and in exposed communities ^7^ or occupational cohorts ^8^. Epidemiological studies further suggest stronger associations between PFAS exposure and hypertension or cardiovascular disease in men ^47^, with meta-analysis showing PFAS-associated risk elevation in men but not in women^23^. Our findings parallel these patterns. That is, male mice accumulated higher PFOA and PFHxS levels than females given the same exposure, and correspondingly developed greater pressor responses, SSBP, and kidney injury. Thus, PFAS-exposed 129S6 mice not only provide a novel model of salt-sensitive hypertension but also mimic sex differences in humans.

Sympathetic nerve activation raises blood pressure by increasing heart rate, peripheral vascular resistance, adrenal catecholamine release, and renal sodium reabsorption ^48^. In our study, the upper PFAS dose increased heart rate in both sexes, but tachycardia neither predicted nor explained blood pressure elevation. That is, it was insufficient to raise blood pressure in females and unnecessary in males. Plasma norepinephrine was unchanged, further arguing against systemic sympathetic activation. Nevertheless, localized sympathetic modulation of renal tubular function remains plausible, as renal efferent nerves directly enhance sodium and water reabsorption, effects that are attenuated by renal denervation ^49,50^. Thus, while systemic sympathoexcitation is unlikely to mediate PFAS-induced hypertension, regional neural control of the kidney merits further study.

The collecting duct plays a critical role in blood pressure regulation through ENaC-mediated fine-tuning of sodium-water balance in the final stages of urine formation ^51^. ENaC abundance is classically regulated by the aldosterone-mineralocorticoid receptor (MR)-SGK1 pathway ^41^. Aldosterone activates MR, inducing the expression of SGK1, which phosphorylates and inhibits Nedd4-2, thereby preventing ENaC ubiquitination/degradation and enhancing its cell surface abundance ^40^. In the present study, αENaC protein was increased by PFAS exposure, but SGK1 protein and phosphorylation were unchanged, arguing against a role of MR-dependent SGK1 expression. While decreased Nedd4-2 Ser448 phosphorylation would typically be associated with an increase of ENaC at the plasma membrane ^40–42^, both ENaC and Nedd4-2 are regulated by many other stimuli including insulin, vasopressin, and other kinases including the metabolic sensor AMP-activated protein kinase (AMPK) ^52^. Our scRNAseq analyses showed enrichment of mitochondrial respiration and energy production pathways across multiple nephron segments following PFAS exposure, supporting the emerging concept that shifts in renal metabolism act as adaptive responses to environmental stressors to sustain tubular transport ^53^. Since AMPK regulates ENaC activity through phosphorylation of Nedd4-2 and AMPK activity alters in response to metabolic changes ^54^, and it is highly attractive to investigate the link between metabolic dysregulation and renal sodium transport in future studies.

The increase in αENaC mRNA, particularly in the lower PFAS dose where SGK1/Nedd4-2 signaling was unaffected, suggests transcriptional upregulation. ENaC transcription is directly stimulated by peroxisome proliferator activated receptor (PPARγ), and global PPARγ activation by thiazolidinediones (TZDs) produces sodium/fluid retention and edema through ENaC-dependent mechanisms in diabetes and heart failure patients ^55^. These adverse effects are recapitulated in TZD-treated mice and are abolished in collecting duct-specific PPARγ knockout mice ^56,57^. Notably, PFAS are PPARγ agonists ^58^, raising the possibility that they promote salt-sensitive hypertension by enhancing ENaC transcription in the collecting duct. This is consistent with our findings of αENaC upregulation in PFAS-exposed mice and the effectiveness of amiloride in normalizing blood pressure during high salt intake. Future studies testing collecting duct-specific PPARγ deletion will be critical to establish this pathway as a mediator of PFAS-induced salt-sensitive hypertension.

While our mechanistic studies focused on ENaC, the contribution of other nephron segments cannot be ruled out. The proximal tubule reabsorbs ∼65% of filtered sodium, primarily via NHE3, but also through sodium-coupled cotransporters such as sodium-glucose cotransporter 2 (SGLT2, *Slc5a2*) and sodium-phosphate co-transporter 2 (Npt2a, *Slc34a1*) ^51^. In our scRNAseq analyses, upper-dose PFAS exposure increased the expression of both SGLT2 and Npt2a (Supplemental Figure 4H-4I). As amiloride only partially corrected salt-induced hypertension in the upper dose, these transporters may have contributed to PFAS-induced pressor effects. In addition, we observed a 50% increase in pNCC/NCC ratio in the lower dose group (Supplemental Figure 6). The phosphorylated Thr53 residue of mouse NCC, orthologous to the stimulatory Thr55 in human NCC, is regulated by the ste20-related proline-alanine-rich kinase (SPAK) and oxidative stress response kinase (OSR1) signaling, suggesting that PFAS may also augment distal convoluted tubule sodium reabsorption through this pathway ^59^. Taken together, our results demonstrate that PFAS exert broad effects on renal sodium transport, and PFAS-induced salt-sensitive hypertension likely reflects the combined actions of multiple mechanisms.

Lastly, the upper PFAS dose induced significant systemic effects, including reduced body weight, striking hepatomegaly, and decreased spleen weight and kidney immune cell content, consistent with prior reports ^33,34^. The increase in liver mass may be mediated by hepatic lipid accumulation and is associated with increased blood lipoprotein cholesterol^60,61^. Interestingly, our scRNAseq analysis of the proximal tubule cells revealed a PFAS-induced expression of low density lipoprotein receptor (LDLR), which has been shown to mediate pathological cholesterol influx and accumulation in renal tubules of Alport mice ^62^. Furthermore, PFAS also upregulated sterol regulatory element-binding protein (SREBP)-2, a transcription factor that regulates expression of genes involved in cholesterol synthesis, including HMG-CoA synthase 2 (*Hmgcs2*) and HMG-CoA reductase (*Hmgcr*) ^38,39^. HMG-CoA reductase is a rate-limiting enzyme in cholesterol synthesis, while HMG-CoA synthase generates its substrate HMG-CoA from acetyl-CoA and acetoacetyl-CoA. Both genes were upregulated in the proximal tubule, particularly in the upper dose group (Supplemental Figure 4), suggesting increased intrarenal cholesterol synthesis. Notably, accumulation of renal cholesterol induces increased expression of pro-fibrotic growth hormones, leading to mesangial expansion, proteinuria, glomerulosclerosis, and tubulointerstitial fibrosis in diabetic nephropathy ^38,39^. Collectively, these observations suggest that PFAS may enhance renal cholesterol uptake and synthesis, leading to cholesterol-mediated lipotoxicity, proximal tubular injury, and kidney fibrosis ^63^.

In summary, our study advances a new paradigm in PFAS toxicology by linking chronic exposure to renal sodium transporter activation, intrarenal metabolic reprogramming, and salt-sensitive hypertension. These results highlight the kidney as a key target of PFAS toxicity and suggest that even low-level exposures may be clinically relevant. Future studies using cell type–specific genetic models and human translational cohorts will be critical to define precise mechanisms and to identify therapeutic strategies to mitigate PFAS-induced cardiovascular and renal risk.

## Data Sharing Statement

The data underlying this article will be shared upon reasonable request to the corresponding authors. The raw data in scRNAseq has been uploaded to the NCBI Sequence Read Archive (SRA) database (ID PRJNA1328035) and will be released on October 1, 2026.

## Acknowledgements

We are grateful to the Histology, Biochemistry, and Molecular Imaging Core at the Center for Musculoskeletal Research at the University of Rochester (supported by NIAMS P30AR069655 grant), and especially Mr. Jeffrey Fox, for assistance with imaging and quantification of mouse renal histology. The single cell RNA sequencing work described in this publication benefited from the support of the Genomics Shared Resource at the Wilmot Cancer Institute, supported in part by the University of Rochester Wilmot Cancer Institute Support Grant #P30CA272302. We thank Dr. Jeffrey Malik, technical director at genomic research center, and the staff members for their technical assistance. The content is solely the responsibility of the authors and does not necessarily represent the official views of the National Institutes of Health.

## Author Contributions

JL, CAK, and VG conducted physiological and molecular biology experiments and analyzed data; QQ and JH performed scRNAseq analyses and interpreted results; CMP and BPL provided the PFAS compounds and designed the quaternary exposure protocol; NLDP and CAM performed mass spectrometry quantitation of PFAS; JL, CAK, CAM, THL, CL, ML, and JOW contributed to the conception of studies, experimental design, and data interpretation. JL drafted the manuscript. QQ, CAK, NLDP and JOW contributed to the drafting of the manuscript. JOW and CAM obtained funding supporting these studies. All authors edited and approved the manuscript.

