## Supplemental Material for "Per- and Polyfluoroalkyl Substances Induces Salt-Sensitive Hypertension by Upregulating Epithelial Sodium Channel - The First Experimental Evidence Supporting Causality"

### METHODS

**Animals and treatments** All protocols were approved by the Animal Care and Use Committees at the University of Rochester and the University of Arizona. Mouse husbandry and care in the study followed the National Institutes of Health (NIH) guidelines. The 129S6 mice were purchased from Taconic Biosciences (129S6/SvEvTac). Mice were maintained on a 0.4% low salt diet (Dyet #113755 AIN-76A purified rodent diet) upon arrival until they were provided with the 4% high salt diet (Dyet #113756 AIN-76A purified rodent diet). Each cage was supplied with environmental enrichments, and the animal facility was maintained on standard 12-hour light-dark cycles. Tap water supplied by the animal housing facilities was provided to mice until they received special water containing the lower and upper concentrations of the PFAS mixture. Mice were randomized into control, lower PFAS dose, or upper PFAS dose groups using a random number sequence generated with the RANDBETWEEN function in Excel. Each mouse was assigned a random number, sorted in ascending order, and then allocated sequentially to the three groups.

The upper dose, consisting of 0.009 mg/mL PFOA + 0.009 mg/mL PFOS + 0.0018 mg/mL PFHxS, and 0.0013 mg/mL PFNA in drinking water [1], which delivered an estimated daily dose of 1.88 mg/kg PFOA, 1.88 mg/kg PFOS, 0.376 mg/kg PFHxS, and 0.26 mg/kg PFNA (Table 1A). The lower dose group received water containing 1:100 dilution of the upper dose, which is estimated to deliver 18.8 µg/kg/day PFOA, 18.8 µg/kg/day PFOS, 3.76 µg/kg/day PFHxS, and 2.6 µg/kg/day PFNA. These estimated daily doses were based on our prior observations that a 20-gram mouse typically consumed 4 mL water/day on a chow diet or a low salt diet [2]. Because water consumption doubled on the 4% high salt diet [2], the concentrations of PFAS added to the water were reduced by 50% in the high salt phase to maintain the same exposure levels. Some mice received amiloride at 20 mg/kg/day in drinking water after salt-induced hypertension was established. Blood pressure and heart rate were chronically monitored using radiotelemetry as previously described [2-4].

**Materials** PFOA (Catalog #171468), PFOS (Catalog #77282), PFHxS (Catalog #50929), and PFNA (Catalog #394459) were purchased from Sigma-Aldrich (St Louis, MO) and were ≥95% pure. Amiloride hydrochloride dihydrate was purchased from Thermo Scientific (Catalog #J62168.06).

**Quantification of PFAS by QTOF-MS** Plasma levels of PFOA, PFOS, PFNA, and PFHxS at the end of the 3-week exposure were measured by liquid chromatography with quadrupole-time-of-flight mass spectrometry (LC-QTOF-MS) on an Agilent Infinity II LC connected to an Agilent 6560 QTOF-MS. All analytical standards including mass-labeled internal standards, injection standards, and native standards, were obtained from Wellington Laboratories. Solvents used throughout the procedure were LC-MS grade and sourced from Fisher Scientific.

Sample preparation and analysis are described in detail in previous studies [1,5] Submandibular bleeding was performed to collect 100-200 µL blood and plasma was snap-frozen in liquid nitrogen, shipped to Carnegie Mellon University, and stored in -80 freezers. On the day of measurement, plasma was thawed to room temperature and processed to remove proteins and lipids using Agilent Enhanced Matrix Removal (EMR) lipid cartridges. Prior to use, the cartridges were pre-conditioned three times by adding 1 mL of 80% acetonitrile (ACN) and 20% Optima LCMS-grade water. Then, a 100 µL aliquot of each plasma sample was added to an EMR lipid cartridge along with 20 µL of a mass-labeled PFAS mixture in ACN (200 ng/mL) and 380 µL of crash solvent (0.1 M formic acid in cold 80% ACN). The mixture was allowed to passively mix for ten minutes. Elution was completed using a positive pressure manifold under ultrahigh purity nitrogen. The eluent was collected in polypropylene vials, to which 20 µL of an injection standard ($^{13}C$-labeled PFOA in ACN, 200 ng/mL) was added. All extracts were stored under refrigeration until analysis. Triplicate blanks were prepared using both Optima water and bovine calf serum (Fisher Scientific, Catalog # 16010159) following the same procedure as the plasma samples to serve as matrix-matched and solvent-based procedural blanks.

A 35 µL volume of each extract was injected onto an Agilent Poroshell C18 column using a binary mobile phase gradient (organic: 100% Optima Methanol and aqueous: 10 mM ammonium acetate in Optima water). The system operated in negative electrospray ionization mode with data-independent acquisition (m/z 65–1100 Da), using three collision energies (0, 10, and 35 eV) for quantification of PFOA, PFOS, PFHxS and PFNA. All analytes were quantified using 1/x weighted linear calibration curves. The linear quantitation range for each analyte was defined using at least five consecutive calibration points with concentrations within ±30% of known values, with an $R^2$ value greater than 0.99 required. The method reporting limit (MRL) for each PFAS was established as the minimum point in the calibration. MRLs (ng/mL) were 1.49 for PFOA, 0.054 for PFOS (linear + branched), 1.151 for PFHxS, and 1.59 for PFNA. A set of triplicate continuing calibration verification serum at 12.5 ng/mL was also analyzed. Recoveries for all compounds in these quality control samples were within ±25% of expected concentrations.

**Radiotelemetry** Blood pressure and heart rate were chronically monitored using radiotelemetry as previously described [2-4]. Briefly, mice at 8 weeks of age were anesthetized with a mixture of ketamine (87.5 mg/kg) and xylazine (12.5 mg/kg) and the pressure catheter was inserted into the left common carotid artery. The transmitter (PA-C10) was subcutaneously placed on the back. Buprenorphine ER (0.6 mg/kg, subcutaneous) was used for peri-operative analgesics. Mice exhibiting excessive pain, suffering or distress that could not be relieved by analgesics were euthanized and excluded from the studies. Mice were allowed to recover for 12 days before measurements were taken on the Ponemah software. Blood pressure and heart rate were recorded for 10 minutes every hour for three consecutive days at baseline and weekly during PFAS exposure protocols. Occasional telemeter catheter clogging occurred during long experimental protocols. After the three-week low-salt diet (Figure 1), clogging was detected in 7 of 55 mice (12%): 3 male controls, 1 male in the lower PFAS group, 2 males in the upper PFAS group, and 1 female in the upper PFAS group. Diagnosis was based on abnormal pressure waveforms, abrupt blood pressure fluctuations, or reduced pulse pressure (<15 mmHg), and confirmed by visual inspection at study termination. These mice did not generate valid telemetry data during the high-salt phase and were excluded from Figure 2.

**Metabolic cage studies and urinary albumin** Additional mice without radiotelemetry were individually housed in metabolic cages (MMC100, Hatteras Instruments) for 48 hours at the end of the high salt diet. Mice had *ad libitum* access to food and water. After acclimation in the first 24 hours, urine was collected in the second 24-hour period. Urinary albumin was measured using an enzyme linked immunosorbent assay (Albuwell M ELISA, Ethos Biosciences, Catalog #1011) following manufacturer's instructions.

**Masson's trichrome staining** Consecutive 6-micron sections were obtained from paraffin-embedded mouse kidneys. For each mouse, two consecutive sections were placed on the same slide and stained with Masson's trichrome blue as previously described [6,7]. Whole slide images were obtained on Olympus VS120 slide scanner at 40x resolution and fibrosis was quantified as the percentage of blue area per 20x field on QuPath version 0.4.2 by a trained technician with no knowledge of experimental groups [8]. Three 20x cortex fields were quantified for each kidney and the average value of the three fields was reported.

**Single-cell RNA Sequencing in whole mouse kidneys** Male 129S6 mice at 8-weeks of age received plain tap water or water containing the lower and upper PFAS doses for 3 weeks on 0.4% low salt diet. Immediately after sacrifice, mice were perfused through the left ventricle with 20 mL heparinized DPBS to remove blood. The kidney was harvested, minced, and incubated in DMEM (Thermo Fisher 11995065) containing 1 mg/mL collagenase A, 1 mg/mL collagenase B, and 25 mM HEPES at 37 °C for 10 min with agitation. Reaction was deactivated by adding 5% FBS and the solution was passed through a 40 μm cell strainer. The single-cell suspension was centrifuged at 500 g for 5 minutes at 4 °C, and the cell pellet was incubated with 3 mL red blood cell lysis buffer on ice for 4 minutes. Cells were centrifuged at 500 g for 5 minutes at °C and the cell pellet were resuspended in 1x DPBS. Cell suspensions were processed to generate single-cell RNA-Seq libraries using Chromium Next GEM Single Cell 3′ GEM, Library and Gel Bead Kit v3.1 (10x Genomics), per the manufacturer's recommendations, as summarized below. First, cell viability and concentration were assessed via trypan staining and automated cell counting using a Biorad TC20. Samples were then loaded on a Chromium Single-Cell Instrument (10x Genomics, Pleasanton, CA, USA) to generate single-cell GEMs (Gel Bead-in-Emulsions). GEM reverse transcription (GEM-RT) was performed to produce a barcoded, full-length cDNA from poly-adenylated mRNA. After incubation, GEMs were broken, the pooled GEM-RT reaction mixtures were recovered, and cDNA was purified with silane magnetic beads (DynaBeads MyOne Silane Beads, ThermoFisher Scientific). The purified cDNA was further amplified by PCR to generate sufficient material for library construction. Enzymatic fragmentation and size selection was used to optimize the cDNA amplicon size and indexed sequencing libraries were constructed by end repair, A-tailing, adaptor ligation, and PCR. Final libraries contain the P5 and P7 priming sites used in Illumina bridge amplification. Libraries were sequenced using Illumina's NovaSeq 6000, targeting 100,000 reads per cell.

Raw fastq files were processed using CellRanger (v7.0.0) to generate feature-barcode matrices based on 10x Genomics mouse reference genome assembly mm10 (refdata-gex-mm10-2020-A). CellBender (v0.2.0) [9] was applied to remove background noise. DoubletFinder R package (v2.0.3) [10] was used to identify and remove potential doublets, with the expected doublet rate set to max(0.3, #cells*8*1e-6).

Further quality control steps included removing cells with fewer than 200 genes, more than 20,000 unique molecular identifiers (UMIs), or a mitochondrial gene content exceeding 30%. For cells passing quality control, the gene count matrix was log-normalized based on total UMI counts and scaled using Seurat R package (v4.3.0.1) [11]. The top 2000 most variable genes were used for principal component analysis (PCA). Batch effects were corrected using Harmony R package (v1.2.0) [12]. Clustering was performed using top 30 principal components with the Louvain algorithm, varying the resolution parameter from 0.1 to 1 in increments of 0.1.

To annotate identified clusters, differentially expressed genes (DEGs) were identified by Wilcoxon rank sum test using FindAllMarkers function in Seurat. Statistical significance was assessed using the Benjamini-Hochberg adjusted false discovery rate (FDR), with a threshold of FDR <0.05. Cluster annotations were guided by reference markers from CellMarker [13] and relevant literature [14,15]. Pathway enrichment analysis for each cell type was performed using fsea R package (v.1.33.1) [16], based on DEGs from the previous step. Gene sets were sourced from the Gene Ontology (GO) database using msigdbr R package (v.7.5.1) [17]. Pathways were considered significantly enriched at an FDR < 0.05. Visualization was performed using ggplot2 (v.3.5.1)[18].

For intra-cell-type comparisons across experimental groups, DEGs were computed using the Wilcoxon rank-sum test using the FindMarkers function in Seurat. Pairwise comparisons included: lower-dose vs. control, upper-dose vs. control, upper-dose vs. lower-dose. DEGs were considered significant if they met both an FDR < 0.05 and an absolute log fold change > 0.25. For a subset of pre-defined genes of interest (e.g., sodium transporters), we applied an alternative filter of FDR < 0.05 and absolute percentage expression difference > 5%, as modest but consistent expression shifts were observed and independently validated by rt-PCR.

**Real-time quantitative PCR** Additional Male 129S6 mice at 8 weeks of age received plain tap water or water containing the lower and upper PFAS doses on low salt for 3 weeks then on high salt for another week. After sacrifice, kidney cortex and medulla were separated under dissecting microscope, snap-frozen in liquid nitrogen, and stored in -80 freezer for real-time quantitative PCR and western blot (see below). To detect the gene expression of different sodium transporters, RNA was extracted from the cortex and medulla. Kidney cortex was processed in ultrasonic homogenization and RNA was extracted using the TRIzol method (Invitrogen, Carlsbad, CA). RNA concentration was measured using a NanoDrop spectrometer with an OD260/OD280 ratio greater than 1.9. Total RNA was reverse transcribed into cDNA using SuperScript III reverse transcriptase (Invitrogen, #18080044) and real-time qPCR was performed using Taqman gene expression assays (Applied Biosystems). The assay numbers for Taqman were follows: Mm01352473_m1 (mouse *Slc9a3*), Mm01275821_m1 (mouse *Slc12a1*), Mm00490213_m1 (mouse *Slc12a3*), Mm00803386_m1 (mouse *Scnn1a*), Mm00441215_m1 (mouse *Scnn1b*), Mm00441228_m1 (mouse *Scnn1g*), and Mm99999915_g1 (mouse *Gapdh*). The individual amount of mRNA was calculated after normalizing to its corresponding internal control using the $2^{-\Delta\Delta Ct}$ method as we previously described [2].

**Western blot** Snap-frozen kidney cortex and medulla samples were thawed on ice and pulse-sonicated in 1x Laemmli sample buffer at a concentration of 20 mg/mL in the presence of protease and phosphatase inhibitors (complete Mini, PhosSTOP, Sigma). Lysates were run on Criterion SDS-PAGE gels and transferred to nitrocellulose membranes. Following transfer, protein loading was confirmed with Ponceau S staining. After removing Ponceau S with Tris-Buffered Saline 0.1% Tween (TBST), membranes were blocked for one hour with 2% BSA in TBST. Primary antibodies were incubated overnight, washed 3x with TBST, then hybridized with HRP-conjugated secondary antibodies (1:5000). Primary antibody information is given in Supplemental Table 1. Following 3 more TBST washes, proteins were visualized with chemiluminescence on a Bio-Rad imager. Densitometry was determined using Image Lab software (Bio-Rad). Proteins were normalized to loading control and then to averaged control value to demonstrate relative changes in expression.

**Statistics** All results were expressed as mean ± SEM. GraphPad Prism (version 10.4) was used for statistical analyses. Time course data of blood pressure and heart rate were analyzed by two-way ANOVA with repeated measurements. Group means were analyzed with one-way ANOVA. Tukey's multiple comparison tests were performed for pairwise comparisons. When standard deviations were significantly different among groups (Brown-Forsythe test and Bartlett's test), the data were analyzed with the Brown-Forsythe ANOVA or Welch ANOVA test. For data not exhibiting Gaussian distribution, Kruskal-Willis test was performed. A p-value <0.05 was considered significant.

**Supplemental Table 1 Antibodies used in western blot.**

| Antibody | Company | Catalog # | Concentration |
|---|---|---|---|
| NCC | StressMarq | SPC-402D | 1:1000 |
| pNCC | PhosphoSolutions | p1311-53 | 1:1000 |
| αENaC | StressMarq | SPC-403D | 1:1000 |
| βENaC | Alomone | ASC-019 | 1:1000 |
| γENaC | StressMarq | SPC-405D | 1:1000 |
| α-$Na^+$/$K^+$ ATPase | Santa Cruz Biotech | sc-21712 | 1:1000 |
| Nedd4-2 | Cell Signaling Tech | 4013S | 1:1000 |
| pNedd4-2 | Cell Signaling Tech | 8063S | 1:1000 |
| SGK1 | Cell Signaling Tech | 12103S | 1:1000 |
| pSGK1 | Cell Signaling Tech | 5599S | 1:1000 |

**Supplemental Table 2 Body weight and organ weight**

| | Male | | | Female | | |
|---|---|---|---|---|---|---|
| | Control | Lower Dose | Upper Dose | Control | Lower Dose | Upper Dose |
| Body Weight (g) | 28.7 ± 1.0 | 27.3 ± 0.5 | 18.3 ± 1.4 * # | 23.9 ± 0.7 | 21.9 ± 0.4 | 20.2 ± 3.1 |
| Heart/BW (mg/g) | 4.6 ± 0.1 | 4.9 ± 0.1 | 6.2 ± 0.2 * # | 5.2 ± 0.2 | 4.5 ± 0.1 | 5.6 ± 0.1 # |
| Spleen/BW (mg/g) | 2.0 ± 0.1 | 2.2 ± 0.1 | 1.6 ± 0.1 * # | 4.6 ± 0.4 † | 3.1 ± 0.1 * | 2.3 ± 0.2 * # |
| Liver/BW (mg/g) | 36.7 ± 1.8 | 38.9 ± 1.6 | 128.0 ± 4.2 * # | 41.3 ± 0.9 | 41.3 ± 1.0 | 134.9 ± 5.1 * # |
| L. Kidney/BW (mg/g) | 6.7 ± 0.3 | 6.9 ± 0.2 | 8.3 ± 0.1 * # | 6.1 ± 0.4 | 5.9 ± 0.1 | 8.4 ± 0.2 * # |
| R. Kidney/BW (mg/g) | 6.6 ± 0.3 | 6.9 ± 0.2 | 8.7 ± 0.1 * # | 6.1 ± 0.3 | 6.3 ± 0.2 | 8.7 ± 0.2 * # |

Body weight and organ weight in mice that completed 3-week low salt followed 1-week high salt diet. Male N = 8/group; Female N = 3 control, 4 lower dose, and 3 upper dose. Data were expressed as mean ± SEM and analyzed with 2-way ANOVA. Body weight: $p_{exposure} < 0.01$, $p_{sex} < 0.05$, $p_{interaction} < 0.05$; Heart/Body weight ratio (mg/g): $p_{exposure} < 0.01$, $p_{sex} = 0.26$, $p_{interaction} < 0.05$; Spleen/Body weight ratio (mg/g): $p_{exposure} < 0.01$, $p_{sex} < 0.01$, $p_{interaction} < 0.01$; Liver/Body weight ratio (mg/g): $p_{exposure} < 0.01$, $p_{sex} = 0.11$, $p_{interaction} = 0.79$; Left Kidney/Body weight ratio (mg/g): $p_{exposure} < 0.01$, $p_{sex} < 0.05$, $p_{interaction} = 0.13$; Right Kidney/Body weight ratio (mg/g): $p_{exposure} < 0.01$, $p_{sex} = 0.08$, $p_{interaction} = 0.37$. Tukey's multiple comparison tests: * p<0.05 vs control; # p<0.05 upper dose vs lower dose; † p<0.05 male vs female at the same exposure level.

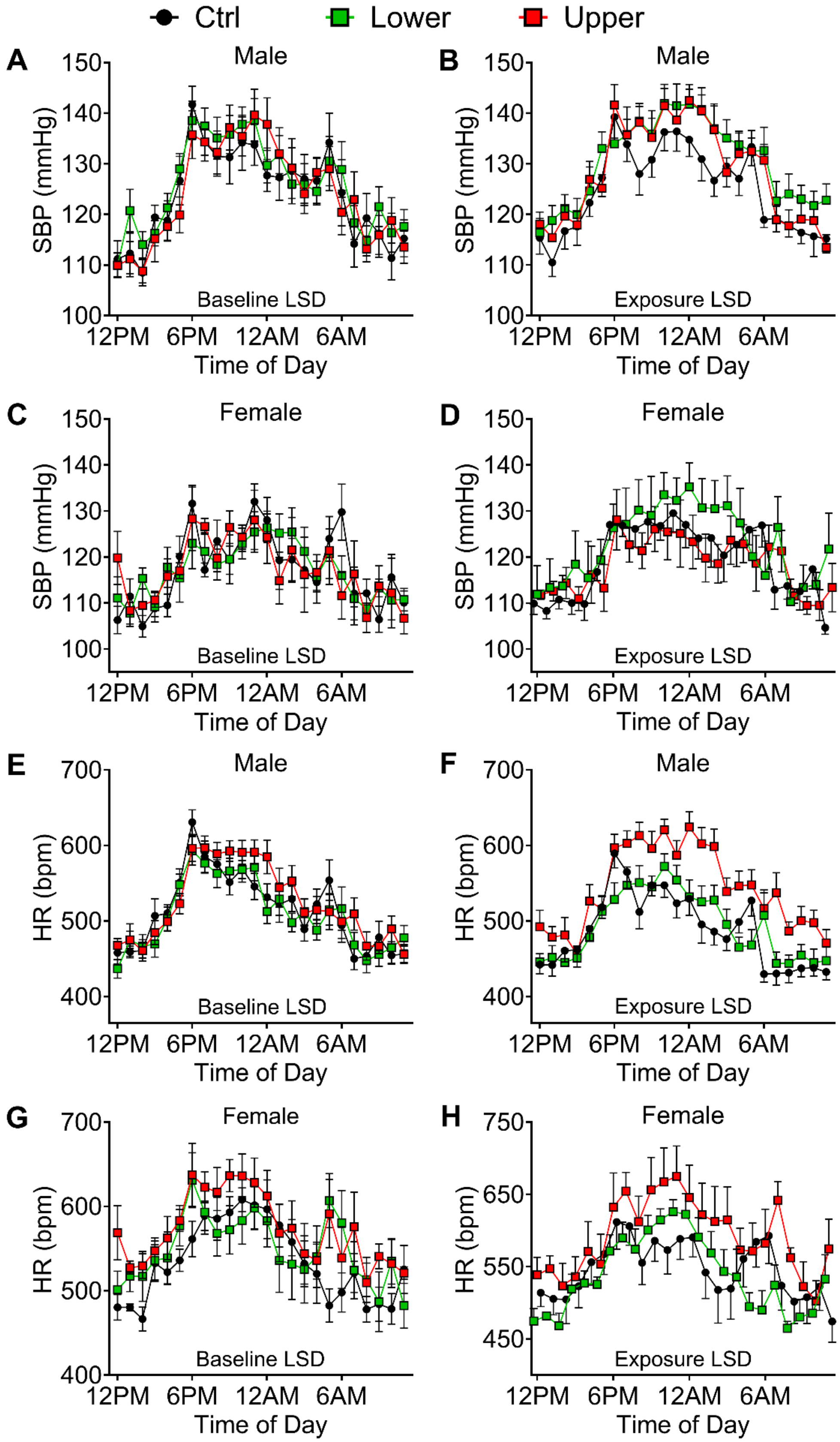

Ctrl
Lower
Upper
A
Male
SBP (mmHg)
150
140
130
120
110
100
Baseline LSD
12PM
6PM
12AM
6AM
Time of Day
B
Male
Exposure LSD
C
Female
D
Female
E
Male
HR (bpm)
700
600
500
400
F
G
Female
H
750
650
550
450

**Supplemental Figure 1 Dose-dependent sex-specific effects of PFAS on blood pressure and heart rate in 129S6 mice on a low salt diet.** Hourly systolic blood pressure (SBP) and heart rate (HR) at baseline and after 3-week exposure were consolidated into 24-hour plots to visualize the circadian rhythm. **A-B**) Male SBP, **C-D**) Female SBP, **E-F**) Male HR, **G-H**) Female HR.

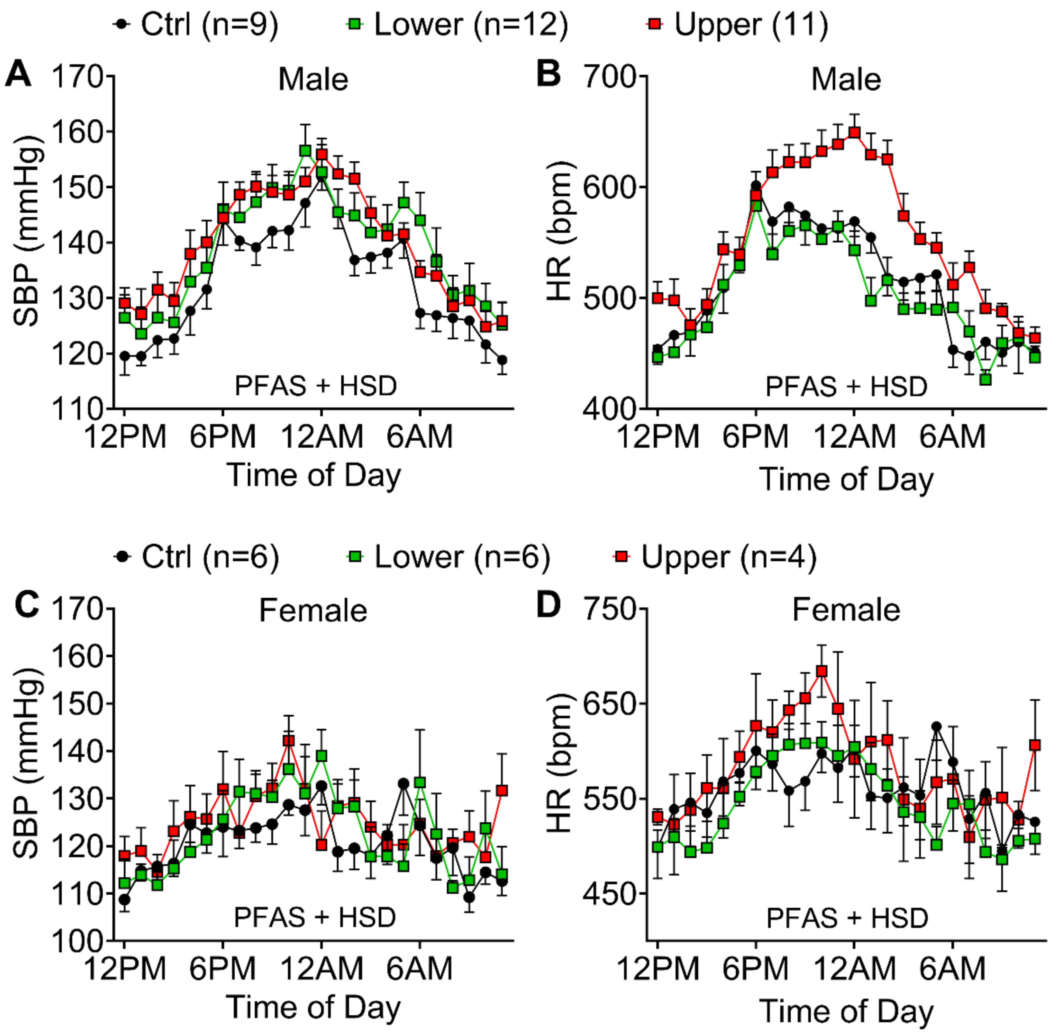


**Supplemental Figure 2 Salt-sensitive hypertension accompanied by tachycardia during exposure to the upper PFAS dose. A-D)** Consolidated 24-hour SBP and HR representing the averages of the 1-week high salt diet in male and female mice (see protocol in Figure 2A). Male, n = 9, 12, 11; Female, n = 6, 6, 4.

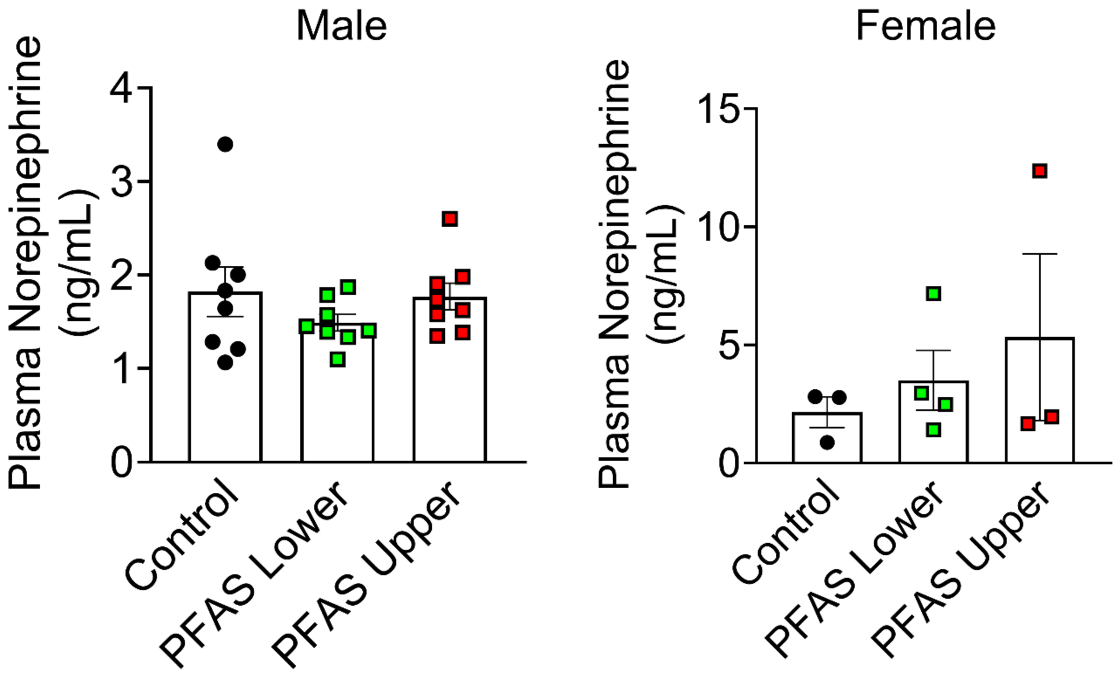


**Supplemental Figure 3 Plasma Norepinephrine Levels.** Blood was collected via cardiac puncture in terminal procedures after the 3-week low salt + 1-week high salt protocol and plasma norepinephrine was determined by ELISA. Data were expressed as mean ± SEM. No statistical significance was detected in one-way ANOVA. Male n = 8; Female n = 3 - 4.

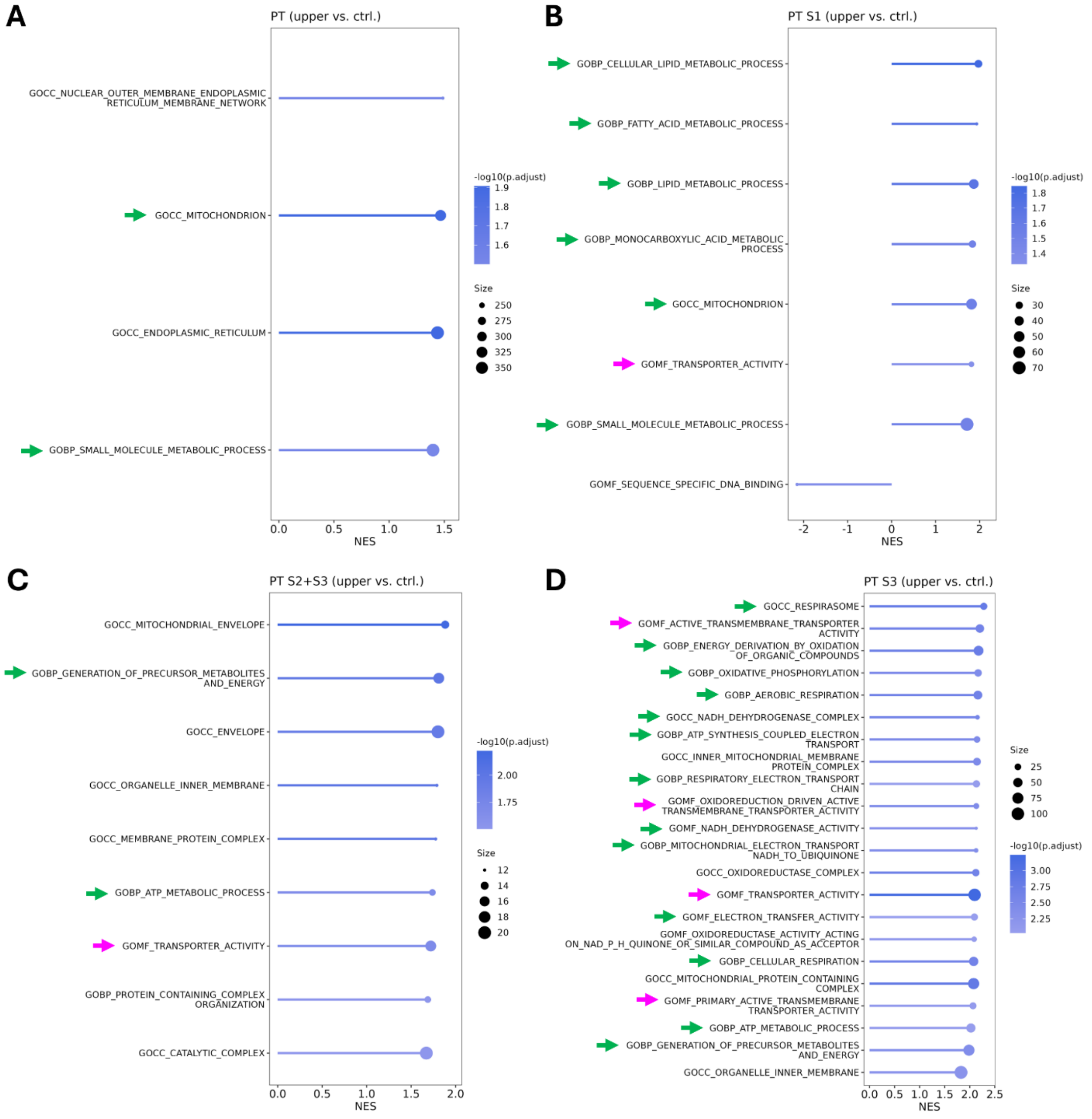

A
PT (upper vs. ctrl.)
GOCC_NUCLEAR_OUTER_MEMBRANE_ENDOPLASMIC_RETICULUM_MEMBRANE_NETWORK
GOCC_MITOCHONDRION
GOCC_ENDOPLASMIC_RETICULUM
GOBP_SMALL_MOLECULE_METABOLIC_PROCESS
NES
-log10(p.adjust)
Size
B
PT S1 (upper vs. ctrl.)
GOBP_CELLULAR_LIPID_METABOLIC_PROCESS
GOBP_FATTY_ACID_METABOLIC_PROCESS
GOBP_LIPID_METABOLIC_PROCESS
GOBP_MONOCARBOXYLIC_ACID_METABOLIC_PROCESS
GOCC_MITOCHONDRION
GOMF_TRANSPORTER_ACTIVITY
GOBP_SMALL_MOLECULE_METABOLIC_PROCESS
GOMF_SEQUENCE_SPECIFIC_DNA_BINDING
NES
-log10(p.adjust)
Size
C
PT S2+S3 (upper vs. ctrl.)
GOCC_MITOCHONDRIAL_ENVELOPE
GOBP_GENERATION_OF_PRECURSOR_METABOLITES_AND_ENERGY
GOCC_ENVELOPE
GOCC_ORGANELLE_INNER_MEMBRANE
GOCC_MEMBRANE_PROTEIN_COMPLEX
GOBP_ATP_METABOLIC_PROCESS
GOMF_TRANSPORTER_ACTIVITY
GOBP_PROTEIN_CONTAINING_COMPLEX_ORGANIZATION
GOCC_CATALYTIC_COMPLEX
NES
-log10(p.adjust)
Size
D
PT S3 (upper vs. ctrl.)
GOCC_RESPIRASOME
GOMF_ACTIVE_TRANSMEMBRANE_TRANSPORTER_ACTIVITY
GOBP_ENERGY_DERIVATION_BY_OXIDATION_OF_ORGANIC_COMPOUNDS
GOBP_OXIDATIVE_PHOSPHORYLATION
GOBP_AEROBIC_RESPIRATION
GOCC_NADH_DEHYDROGENASE_COMPLEX
GOBP_ATP_SYNTHESIS_COUPLED_ELECTRON_TRANSPORT
GOCC_INNER_MITOCHONDRIAL_MEMBRANE_PROTEIN_COMPLEX
GOBP_RESPIRATORY_ELECTRON_TRANSPORT_CHAIN
GOMF_OXIDOREDUCTION_DRIVEN_ACTIVE_TRANSMEMBRANE_TRANSPORTER_ACTIVITY
GOMF_NADH_DEHYDROGENASE_ACTIVITY
GOBP_MITOCHONDRIAL_ELECTRON_TRANSPORT_NADH_TO_UBIQUINONE
GOCC_OXIDOREDUCTASE_COMPLEX
GOMF_TRANSPORTER_ACTIVITY
GOMF_ELECTRON_TRANSFER_ACTIVITY
GOMF_OXIDOREDUCTASE_ACTIVITY_ACTING_ON_NAD_P_H_QUINONE_OR_SIMILAR_COMPOUND_AS_ACCEPTOR
GOBP_CELLULAR_RESPIRATION
GOCC_MITOCHONDRIAL_PROTEIN_CONTAINING_COMPLEX
GOMF_PRIMARY_ACTIVE_TRANSMEMBRANE_TRANSPORTER_ACTIVITY
GOBP_ATP_METABOLIC_PROCESS
GOBP_GENERATION_OF_PRECURSOR_METABOLITES_AND_ENERGY
GOCC_ORGANELLE_INNER_MEMBRANE
NES
Size
-log10(p.adjust)

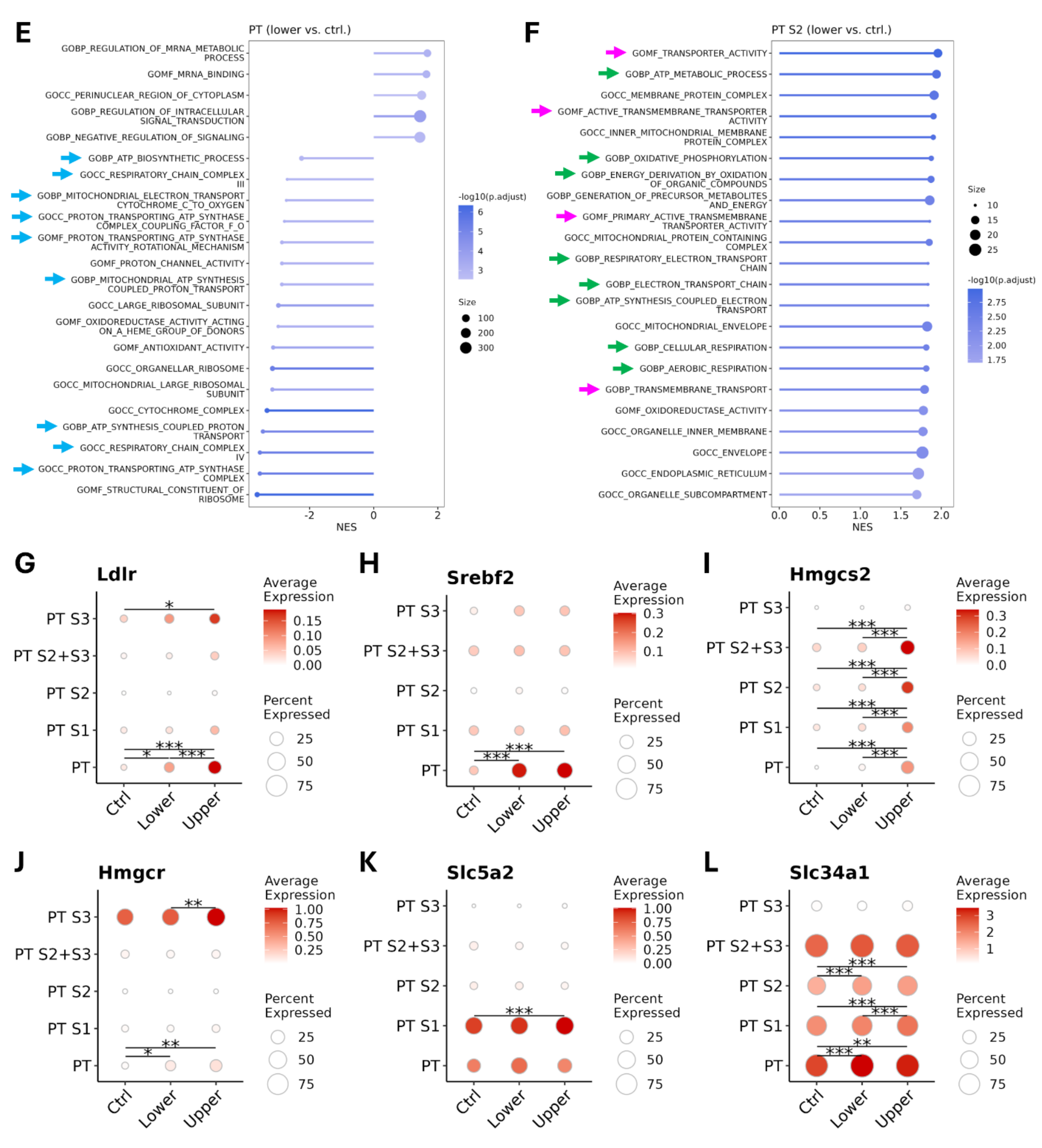


**Supplemental Figure 4 Pathway enrichment analysis and select genes in the proximal tubule. A-D**) Pathway enrichment analysis of DEGs from the PT, PT-S1, PT S2+S3 and PT-S3 in the upper PFAS dose compared to controls. No significantly enriched pathways (FDR<0.5) were identified in PT-S2 segment in response to the upper dose. NES, normalized enrichment score. Positive NES stands for upregulated pathways and negative NES denotes downregulated pathways. Green arrows highlight pathways upregulated in fatty acid metabolism, mitochondria electron transfer chain, oxidative phosphorylation, and energy production. Magenta arrows highlight pathways involved in active transmembrane transporter activities. **E**) Enriched pathways in PT-S2 cells in response to the lower PFAS dose. No significantly enriched pathways were identified in PT-S1, S2+S3, or S3 cells in the lower PFAS dose group. **F**) A PT cell population distinct from S1-S3 segments exhibiting downregulated pathways in mitochondria metabolism and energy production (blue arrows). **G-L**) Dot plots showing cell type-specific expression of select genes across control, lower-dose, upper-dose groups for **G**) *Ldlr*, **H**) *Srebf2*, **I**) *Hmgcs2*, **J**) *Hmgcr*, **K**) *Slc5a2*, **L**) *Slc34a1*. For intra-cell-type comparisons, DEGs were computed using the FindMarkers function in Seurat. ** $p<0.01$, *** $p<0.001$, Wilcoxon rank-sum test.

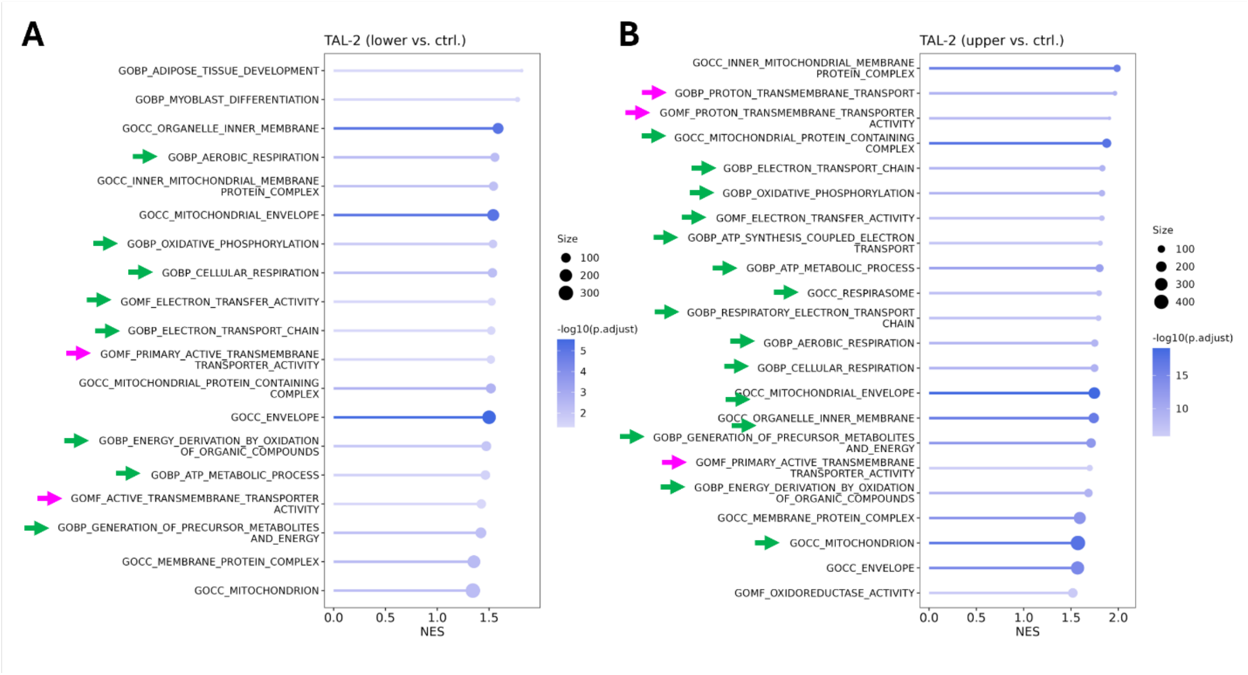


**Supplemental Figure 5 Pathway enrichment analysis of DEGs in thick ascending limb cells**. Significantly enriched pathways (FDR<0.5) were identified in a thick ascending limb cell population (TAL-2, see Figure 4C UMAP plot) in response to the lower and upper PFAS doses. NES, normalized enrichment score. Positive NES stands for upregulated pathways. Green arrows highlight pathways involved in cellular respiration, mitochondria electron transfer chain, oxidative phosphorylation, and energy production. Magenta arrows highlight pathways involved in active transmembrane transporter activities.

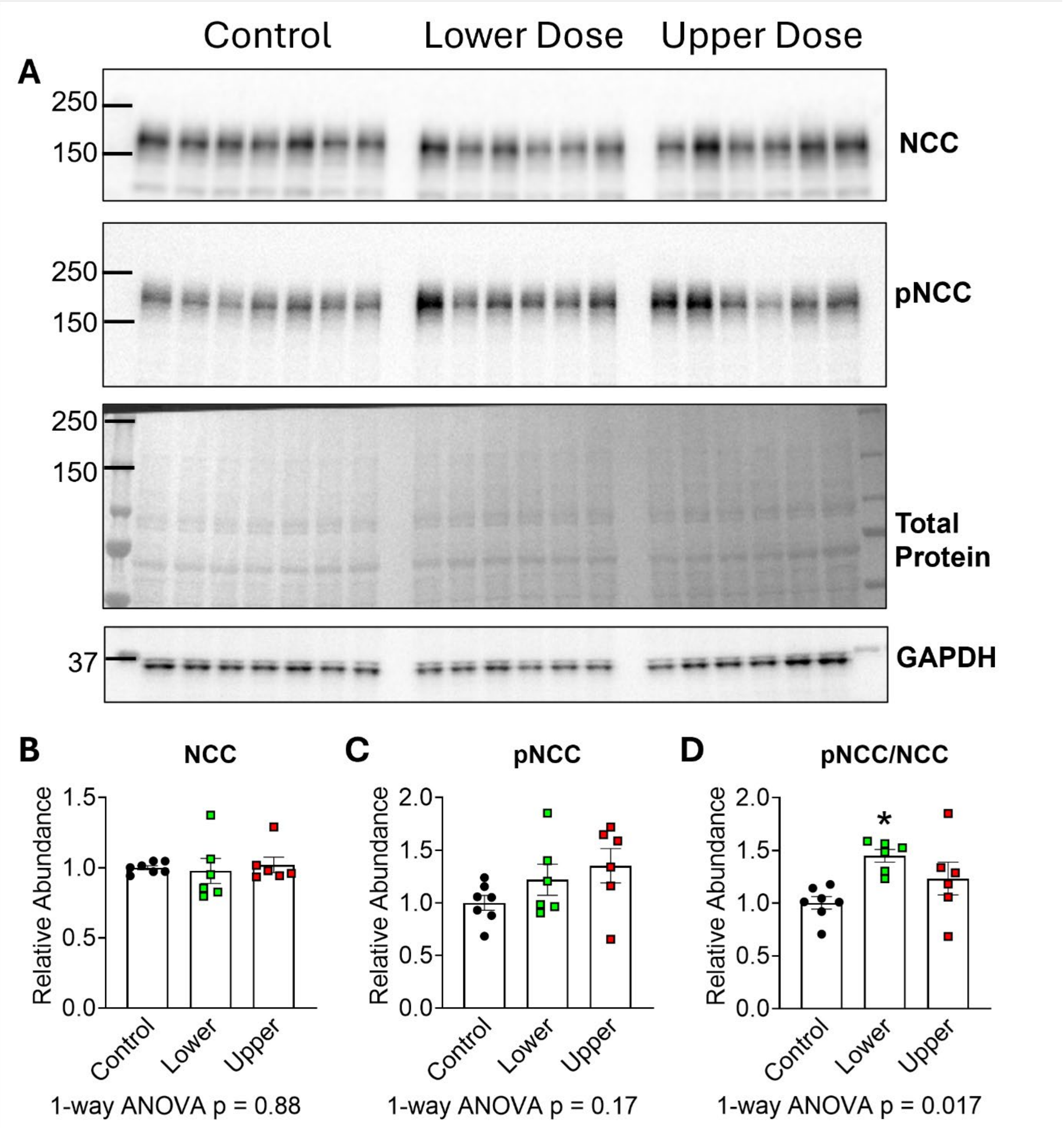


**Supplemental Figure 6 Effects of PFAS exposure on NCC expression and phosphorylation. A)** Control and PFAS-exposed male 129S6 mice received the 0.4% low salt diet for 3 weeks followed by the 4% high salt diet for a week and kidney cortex homogenates were used for western blot. Each lane/dot is an individual mouse; N = 7 control, 6 lower dose, and 6 upper dose. **B-D**) Densitometry for NCC, pNCC (Thr53), and pNCC/NCC ratio. Data expressed as mean ± SEM. One-way ANOVA with Dunnett's post hoc analysis. * $p < 0.05$ compared to control.

### Supplemental Material Reference
